# Hypoimmunogenic hPSC-derived cardiac organoids for immune evasion and heart repair

**DOI:** 10.1101/2025.04.09.648007

**Authors:** Sophia E. Silver, Alessandro R. Howells, Dimitrios C. Arhontoulis, Lauren N. Randolph, Nathaniel A. Hyams, Ryan W. Barrs, Mei Li, Charles M. Kerr, Rob A. Robino, Jordan E. Morningstar, Jacelyn D. Bain, Martha E. Floy, Russell A. Norris, Xiaoping Bao, Jean Marie Ruddy, Sean P. Palecek, Leonardo M. R. Ferreira, Xiaojun Lance Lian, Ying Mei

## Abstract

Human pluripotent stem cell (hPSC)-derived cardiac therapies hold great promise for heart regeneration but face major translational barriers due to allogeneic immune rejection. Here, we engineered hypoimmunogenic hPSCs using a two-step CRISPR-Cas9 strategy: (1) B2M knockout, eliminating HLA class I surface expression, and (2) knock-in of HLA-E or HLA-G trimer constructs in the AAVS1 safe harbor locus to confer robust immune evasion. Hypoimmunogenic hPSCs maintained pluripotency, efficiently differentiated into cardiac cell types that resisted both T and NK cell-mediated cytotoxicity *in vitro*, and self-assembled into engineered cardiac organoids. Comprehensive analyses of the hypoimmunogenic cells and organoids revealed preservation of transcriptomic, structural, and functional properties with minimal off-target effects from gene editing. *In vivo*, hypoimmunogenic cardiac organoids restored contractile function in infarcted rat hearts and demonstrated superior graft retention and immune evasion in humanized mice compared to wild-type counterparts. These findings establish the therapeutic potential of hypoimmunogenic hPSC-CMs in the cardiac organoid platform, laying the foundation for off-the-shelf cardiac cell therapies to treat cardiovascular disease, the leading cause of death worldwide.

## INTRODUCTION

Cardiovascular disease is the cause of 1 out of every 4 deaths in the United States, with over 800,000 people suffering from myocardial infarction (MI) each year^1,2^. Due to the limited regenerative capacity of adult human hearts, human pluripotent stem cell derived cardiomyocytes (hPSC-CMs) have emerged as a powerful cell source for cardiac repair^3–8^. hPSC-CMs have been shown to restore the contractile function of infarcted hearts in numerous models, including non-human primates^9–22^. Preliminary data released from ongoing clinical trials have indicated therapeutic potency of hPSC-CMs to treat heart failure patients^23,24^.

Direct intramyocardial injections of hPSC-CMs into infarcted myocardium provides a straightforward method for heart remuscularization. However, this approach has been limited by low cell retention, suboptimal survival and engraftment, and moderate functional improvement^25,26^. To address these challenges, Laflamme and coworkers systematically investigated cell death pathways post-implantion and developed a “pro-survival” cocktail that achieved robust hPSC-CM engraftment in both rodent and primate models^10,12–14,27^. Tissue engineering strategies have also been utilized to enhance hPSC-CM survival and engraftment. For example, prevascularized cardiac tissue patches have been fabricated to prevent anoikis-mediated cell death and improve anastomosis with host vasculature^9,17–19,21,28–33^. Furthermore, injectable spherical microtissues (i.e., spheroids) composed of monocellular hPSC-CMs have been developed to improve cell retention^20,22^. To introduce prevascularization and myocardium-mimetic environments to the microtissues, we have developed engineered human cardiac organoids by assembling hPSC-CMs with vascular-supporting cells^34,35^. This modular approach allows for the incorporation of additional cells and factors to enhance survival, engraftment, and host integration post-implantation^36^.

Beyond survival and engraftment, immune rejection of allogeneic hPSC-CMs post-implantation remains a critical barrier, necessitating long-term immunosuppression that compromises patient health and life quality^37^. Notably, recent findings of immune rejection of autologous PSC-CMs in non-human primates further underscore the need for immune-evasive cardiac cell therapies^24^. A major mechanism of allogeneic immune rejection is human leukocyte antigen (HLA) class I-mediated CD8^+^ T cell cytotoxicity. While knocking out (KO) Beta-2 Microglobulin (B2M), which encodes a subunit essential for HLA class I surface expression, mitigates T cell-mediated rejection, it simultaneously triggers natural killer (NK) cell-mediated lysis due to the absence of self-recognition signals^38^. To overcome this, NK cell inhibitory molecules, such as CD47 and HLA-E, and HLA-G, have been introduced into B2M^-/-^ cells to generate hypoimmunogenic cells^39–42^ (**Fig. 1a**). This strategy has been successfully implemented to create hypoimmune hPSC-derived beta cells and primary pancreatic islets with long term survival in immunocompetent, allogeneic rodents and primates to treat type 1 diabetes^43–45^. However, the application of hypoimmune engineering to hPSC-CMs for heart repair remains largely unexplored.

**Fig. 1:**
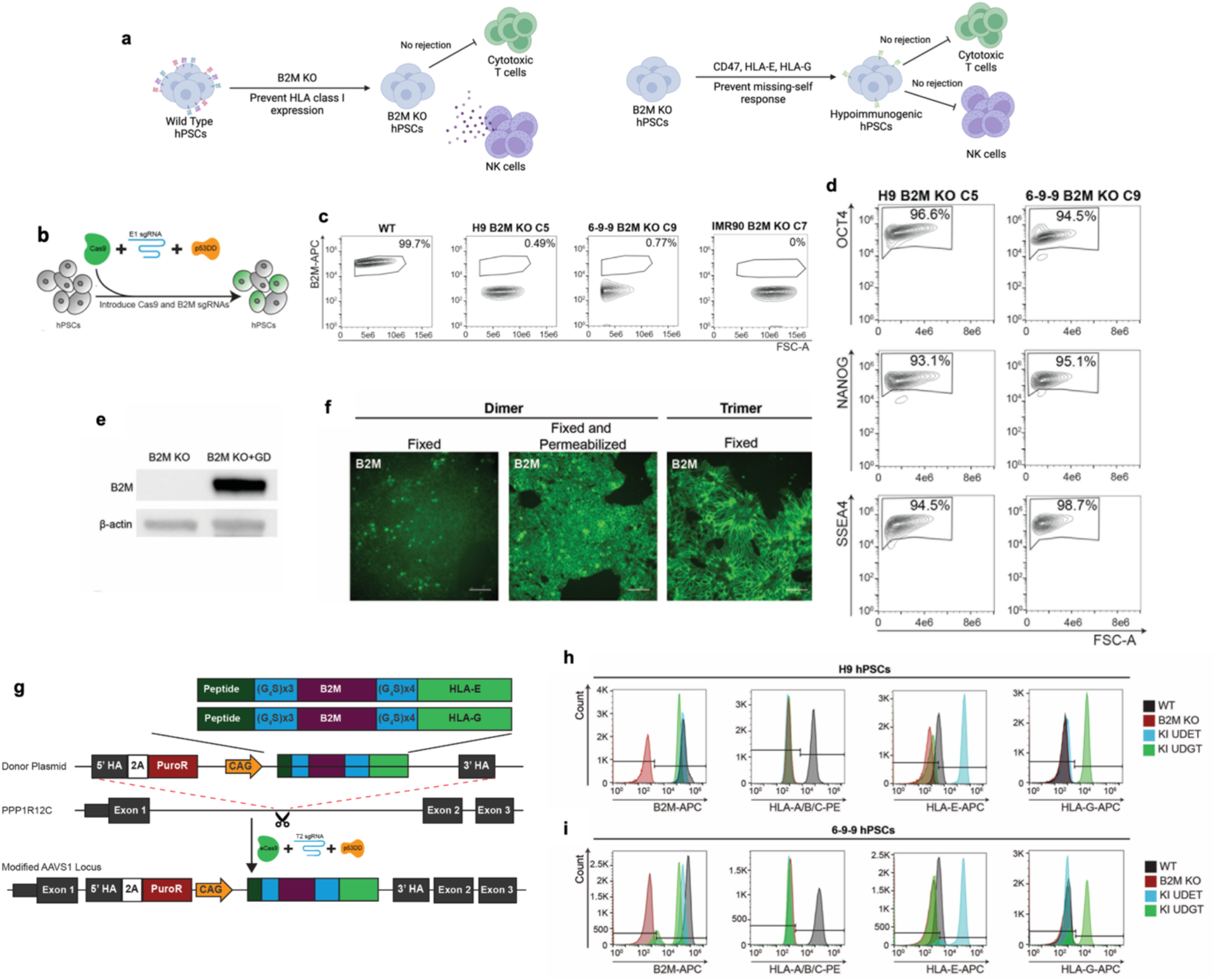
Generation of universal donor hPSCs via biallelic B2M KO and genetically stable KI of non-classical HLA class I trimer constructs. a, Schematic illustrating general engineering strategies previously used to generate hypoimmunogenic hPSCs. b, Experimental schematic for the generation of B2M KO hPSCs, illustrating plasmids encoding Cas9 and GFP, E1 gRNA, and p53DD. c, Flow cytometry analysis of B2M expression for WT and selected single cell derived clones for H9, 6-9-9, and IMR90. d, Flow cytometry analysis of B2M KO hPSC expression of pluripotency markers. e, Western blot results for B2M KO cells and B2M KO+GD cells probed for B2M. ß-actin was used as a housekeeping protein. f, Immunofluorescent microscopy analysis of B2M expression in B2M KO+GD cells (fixed or fixed and permeabilized) and UDGT cells (fixed). Scale bars are 100 µm. g, Schematic of our knock in design and experimental execution, illustrating plasmids encoding eCas9, T2 sqRNA, our donor construct, and p53DD. h-i, Flow cytometry analysis of HLA expression in WT (black), B2M KO (red), KI UDET (blue), and KI UDGT (green) for single cell derived clones for (h) H9 and (i) 6-9-9 hPSCs.

Here, we combined CRISPR-Cas9 genome editing with engineered organoid technology to develop hypoimmunogenic hPSC-derived cardiac organoids for heart repair, knocking out *B2M* and knocking in HLA-E or HLA-G trimer constructs to generate hypoimmunogenic hPSCs. B2M^-/-^ HLA-E^+^ hPSCs were then differentiated into cardiomyocytes (hPSC-CMs), cardiac fibroblasts (hPSC-cFbs), endothelial cells (hPSC-ECs), and pericytes (hPSC-PCs), which then self-assembled into engineered cardiac organoids. RNA sequencing revealed minimal transcriptomic differences between hypoimmunogenic and wild-type (WT) cells and organoids, except for differential HLA-E expression. Structural and functional assessments further supported the similarity between hypoimmunogenic and WT cells/organoids. In a rat model of ischemia/reperfusion injury, hypoimmunogenic cardiac organoids restored contractile function comparably to WT organoids Strikingly, hypoimmunogenic cardiac organoids demonstrated markedly superior graft retention and immune evasion in humanized NSG mice compared to their WT counterparts. For the first time, our study demonstrates that hypoimmunogenic gene editing preserves the therapeutic efficacy of cardiac cell therapy while uniquely enabling immune evasion in the setting of a functional human allogeneic immune system.

## RESULTS

### Design and characterization of B2M KO hPSCs

To inform the design of our universal donor cells for cardiac regenerative therapy, we first examined the transcriptomic HLA expression profile of mature hPSC-CMs, which revealed high expression of classical class I and no class II HLA expression (**Supplemental Fig. 1a**). Our universal donor stem cell design involved immunoengineering HLA expression in two parts, inspired by the structure of class I HLA molecules (**Supplemental Fig. 1b**). We first removed all class I HLA expression by CRISPR-mediated biallelic KO of beta-2-microglobulin (B2M). We designed two single guide RNAs (sgRNAs) (**Supplementary** Fig. 1c), one targeting B2M exon 1 (E1) and one targeting B2M exon 2 (E2) and tested their efficacy alone and in combination in HEK293 cells (**Supplementary** Fig. 1d). We found all three conditions resulted in HEK293 B2M negative cells (**Supplementary** Fig. 1e) and selected our E1 sgRNA for use in hPSCs due to its target location near the start codon. We transfected three separate hPSC lines in our first generation of B2M KO hPSCs (**Fig. 1b**). The inclusion of p53DD in our system greatly improved cell survival following transfection (**Supplementary** Fig. 2a) with a transfection and KO efficiency of 38% GFP+ and 10%, respectively (**Supplementary** Fig. 2b,c). Following single-cell derived colony selection, we identified multiple KO clones in each cell line (**Fig. 1c and Supplementary** Fig. 2d-f), confirmed maintenance of pluripotency (**Fig. 1d**), and validated differentiation potential into each of the three germ layers of the selected B2M KO hPSCs (**Supplementary** Fig. 2g,h). Allelic sequencing revealed 6-9-9 cells with different insertion/deletion patterns for each allele, while H9 cells had the same 10-bp deletion and IMR90 cells had the same 1-bp deletion on both alleles (**Supplementary** Fig. 2i). Together, these data illustrate the successful generation of B2M KO hPSCs clonal lines via CRISPR-Cas9.

### Design and characterization of universal donor hPSCs

The second step in creating universal donor cells involved ensuring expression of nonclassical HLA class I molecules on the cell surface to suppress NK cell and T cell activation, namely the non-polymorphic class I molecules HLA-E and HLA-G^46–52^. We selected a membrane-bound HLA-G isoform for our first fusion protein as it retains all three alpha domains, complexes with B2M, and has been implicated in immune tolerance (**Supplementary** Fig. 3a)^53–55^. We compared surface expression of a synthetic dimer containing B2M and HLA-G (B2M-HLA-G Dimer; “GD”) **(Supplementary** Fig. 3b) and a HLA-G trimer (peptide-B2M-HLA-G Trimer) (**Supplementary** Fig. 3c), which includes a peptide sequence most commonly presented by HLA-G at the maternal-fetal interface^56^. Western blot analysis of B2M suggested GD constructs were successfully expressed in cells (**Fig. 1e**). However, immunostaining and flow cytometry revealed minimal surface expression of B2M in B2M KO+GD cells, whereas B2M KO cells transduced with HLA-G trimer construct (Universal Donor HLA-G Trimer; “UDGT”) showed strong surface expression of B2M, indicating the importance of including the presenting peptide within the construct design to ensure robust surface transport (**Fig. 1f and Supplementary** Fig. 3d). We evaluated the efficacy of our engineered HLA expression and found that WT cells showed expression of B2M and HLA-A/B/C without HLA-E or HLA-G, whereas B2M KO cells showed neither B2M nor HLA expression, and UDGT cell did not show expression of HLA-A/B/C, but showed both B2M and HLA-G expression, independent of IFNγ treatment (**Supplementary** Fig. 3e,f).

In engineering HLA-E expression, we selected a peptide derived from the HLA-C signaling peptide sequence (**Supplementary** Fig. 3g)^51^. B2M KO cells transduced with HLA-E trimer (Universal Donor HLA-E Trimer; “UDET”) also resulted in cells with no expression of HLA-A/B/C, but strong surface expression of B2M and HLA-E, independent of IFNγ exposure (**Supplementary** Fig. 3h,i). Taken together, these data indicate that B2M-HLA dimer design fail to enable surface expression of HLA-G, whereas engineered HLA-E and HLA-G trimer surface expression was achieved through fusing a peptide, B2M and HLA-E/G within the construct design.

### Genetically stable knockin (KI) of HLA trimer constructs

We initially used lentiviral delivery to introduce the UDGT and UDET constructs into B2M KO hPSCs. However, in cardiac progenitor cells (hPSC-CPCs) derived from UDET and UDGT hPSCs, we observed reduced expression of HLA-E and HLA-G (**Supplementary** Fig. 4a,b), likely due to epigenetic silencing during cardiac differentiation. To address this limitation, we modified our integration strategy by employing CRISPR-Cas9–mediated knock-in at the AAVS1 safe harbor locus(**Fig. 1g**)^57–59^. Using this strategy, we achieved approximately 2% KI efficiency before drug selection (**Supplementary** Fig. 4c,d). This efficiency was greatly improved after drug selection. We identified heterozygous KI clones from each line, henceforth referred to as “KI UDET” and “KI UDGT” cells. Analysis of HLA expression demonstrated that KI UDGT and KI UDET hPSCs expressed expected HLA profiles independent from IFNγ stimulation (**Fig. 1h,I and Supplementary** Fig. 4e,f) and remained pluripotent (**Supplementary** Fig. 4g,h). We assessed the cardiac differentiation capacity of KI UDET and KI UDGT hPSCs and observed over 90% efficiency in generating hPSC-CPCs (**Supplementary** Fig. 5a,b). Together, these data support the implementation of the AAVS1 locus as a robust safe harbor site, preserving the differentiation capacity of KI cells and promoting stable surface expression of HLA-E and HLA-G, independent of IFNγ stimulation.

### *In vitro* and *in vivo* analysis of KI UDET hPSC-CPC immunogenicity

Next, we assessed the ability of universal donor hPSC-CPC derivatives to evade immune rejection as a proof of principle to develop hypoimmunogenic cardiac cell-based therapies. We first confirmed that hPSC-CPCs maintained expected HLA expression profiles, with B2M KO hPSC-CPCs showing no HLA/B2M surface expression, WT hPSC-CPCs demonstrating a high dependence on IFNγ to upregulate expression of HLA -A/B/C, and KI UDET hPSC-CPCs showing stable surface expression of HLA-E only, independent of IFNγ treatment (**Fig. 2a**). To assess the efficacy of hypoimmunogenic gene edits in hPSC-CPCs, we performed *in vitro* luciferase release assays using luciferase-expressing hPSC-CPCs and either human donor CD8^+^ T cells or NK cells as target and effector cells, respectively^44^. As expected, when cultured with CD8^+^ T cells, WT hPSC-CPCs were lysed while B2M KO cells were completely protected. We observed statistically significant decrease in lysis of KI UDET and KI UDGT cells compared to WT cells, demonstrating the retained protective effect of their B2M KO (**Fig. 2b**). When cultured with NK cells, we observed an expected inverse trend for WT and B2M KO cell lysis: WT cells evaded detection and B2M KO cells were readily lysed (**Fig. 2c**). KI UDET cells demonstrated a significant reduction in NK cell mediated lysis compared to B2M KO cells, such that there was no significant difference between the KI UDET, KI UDGT, and WT cells. (**Fig. 2c**).

**Fig. 2:**
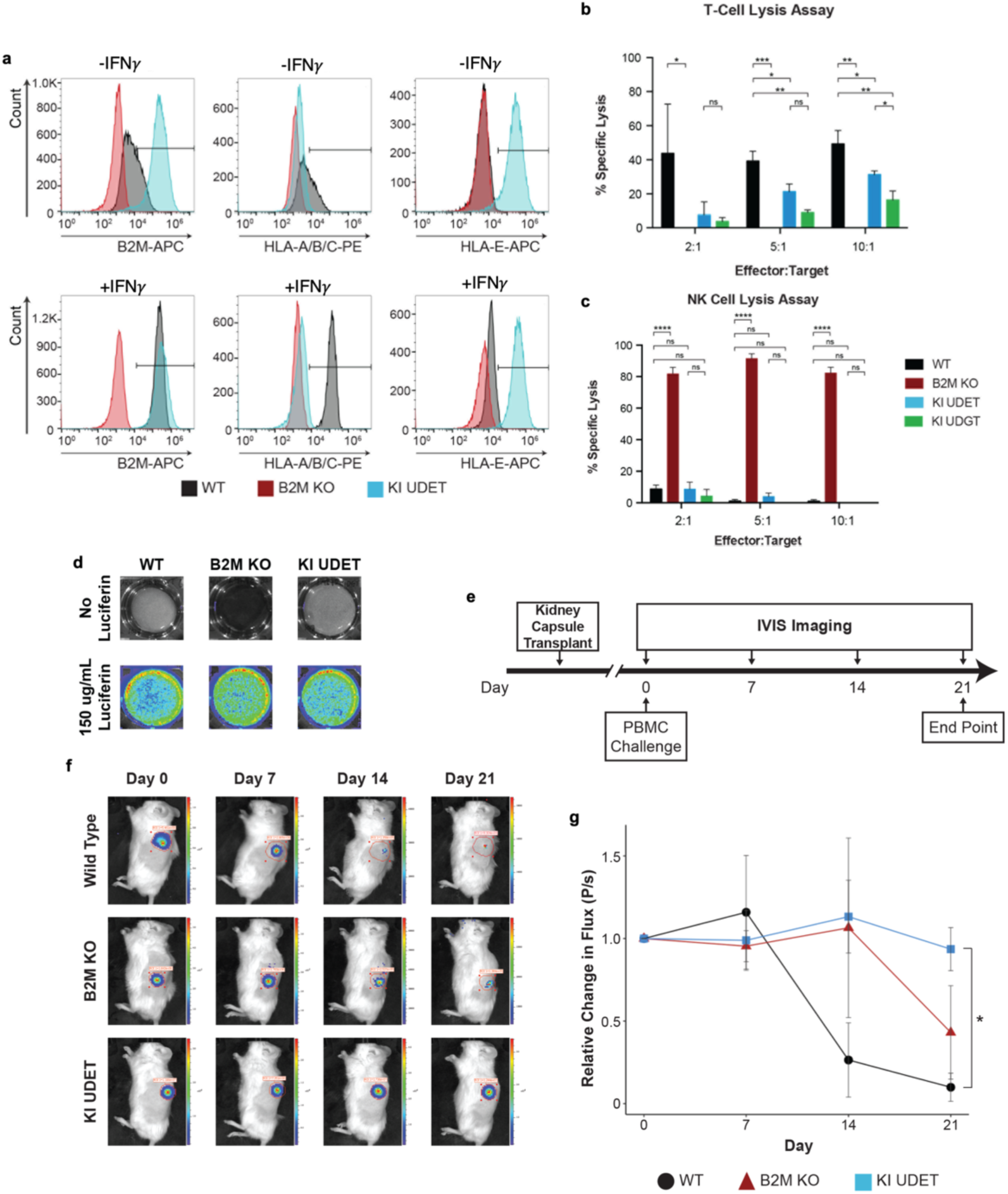
*In vitro* and *in vivo* immunogenicity of universal donor hPSC-CPCs. a, Flow cytometry analysis of HLA expression in day 20 CPCs derived from IMR90 WT (black), B2M KO (red), and KI UDET (blue) lines, treated either with (Bottom) or without (Top) INFγ for 3 days. **b-c**, (**b**) CD8+ T-cell and (**c**) NK cell lysis of hPSC-CPCs measured as percent specific lysis. Results for positive control hPSC-CPCs treated with Triton-X and negative control untreated hPSC-CPCs are incorporated into the calculation of percent specific lysis. **d**, IVIS images of *in vitro* undifferentiated IMR90 GAPDH-Luc hPSCs, treated either with (Bottom) or without (Top) Luciferin. **e**, Schematic of humanized NSG mouse PBMC immune challenge. **f**, Representative IVIS images of WT, B2M KO, and KI UDET mice during each time point analysis, with the region of interest depicted. **g,** Quantification of average change in flux, relative to day 0, as a function of time, with mice that received transplantation of WT (black), B2M KO (red), or KI UDET (blue) hPSC-CPCs (n=3). Error bars represent standard error of the mean. *, **, ***, and **** indicate p values less than 0.05, 0.01, 0.001, and 0.0001, respectively. ns indicates values that are not significantly different.

We progressed in assessing immunogenicity of universal donor hPSC-CPCs *in vivo*, utilizing a humanized NSG mouse model transplanted with human peripheral blood mononuclear cells (PBMCs) and luciferase-expressing KI UDET hPSC-CPCs, selected due to the capacity of HLA-E to bind to the inhibiting receptor NKG2A expressed on most NK cells^60^ (**Fig. 2d,e**). Mice that received WT cells showed a dramatic decrease in relative bioluminescence, indicative of mass cell rejection, whereas mice with B2M KO cells had a delayed decrease in signal, likely due to the low NK cell content within our engrafted PBMCs. In contrast, mice with KI UDET hPSC-CPCs showed a relatively constant signal at significantly higher levels to that of WT signal at Day 21, indicating their resistance to allogeneic immune rejection. (**Fig. 2f,g**).

### Differentiation of KI UDET hPSCs into cardiac cell types

Following proof of principle of immune evasion of derivatives of our KI UDET cells, we moved forward with discerning differences between WT and KI UDET hPSC-derived products, specifically from 19-9-11 hiPSCs. We validated that 19-9-11 KI UDET hPSCs were positive for pluripotency markers (**Supplementary** Fig. 6a-d) and displayed a normal karyotype ∼30 passages after hypoimmunogenic editing (**Supplementary** Fig. 6e). We also confirmed that KI UDET hPSCs from the 19-9-11 line demonstrated less than 1% expression of HLA-A2 (HLA-A allele specific to 19-9-11 cells) and HLA-B/C as well as over 95% HLA-E expression (**Supplementary** Fig. 6f-i). Comparison of whole-transcriptome using principal component analysis (PCA) showed that PC1 accounted for 99% of the explained variance between WT and KI UDET hPSCs. Importantly, HLA-E was the top gene driving PC1 loadings (**Supplementary** Fig. 6j,k) and normalized expression levels of HLA-E were significantly increased in KI UDET hPSCs (**Supplementary** Fig. 6l).

We next sought to differentiate WT and KI UDET hPSCs into cardiac cell types to assess their differentiation capacity and equivalence. We confirmed that both WT and KI UDET hPSCs robustly differentiated into hPSC-CMs, hPSC-cFbs, hPSC-ECs, and hPSC-PCs (**Fig. 3a and Supplementary** Fig. 7a-d) via their expression of cell type-specific markers (**Fig. 3b-m**). Bulk RNA-sequencing analysis indicated distinct clustering between WT and KI UDET hPSC-derived cardiac cells (**Supplementary** Fig. 8a-d). PCA revealed that PC1 accounted for over 99% of the explained variance between WT and KI UDET hPSC-derived cardiac cell types, with HLA-E remaining a primary driver (**Fig. 3n-q and Supplementary** Fig. 8e-h). Differential expression analysis (DEA) further elucidated differences between hPSC-derived cell types, with a focus on genes involved in cardiac and vascular development, extracellular matrix (ECM) production, and immune processes (**Supplementary** Fig. 8i). HLA-E consistently showed significantly higher normalized expression in KI UDET over WT hPSC-derived cells, while DEA revealed no significant alteration to key functional gene sets for each hPSC-derived cell type (**Fig. 3r-u** and **Supplementary** Fig. 8i-m). Together, phenotypic characterization and transcriptomic analysis demonstrate that WT and KI UDET hPSCs and their cardiac cell derivatives display only subtle differences and are nearly transcriptomically identical, excluding variation in the expression of HLA-E.

**Fig. 3:**
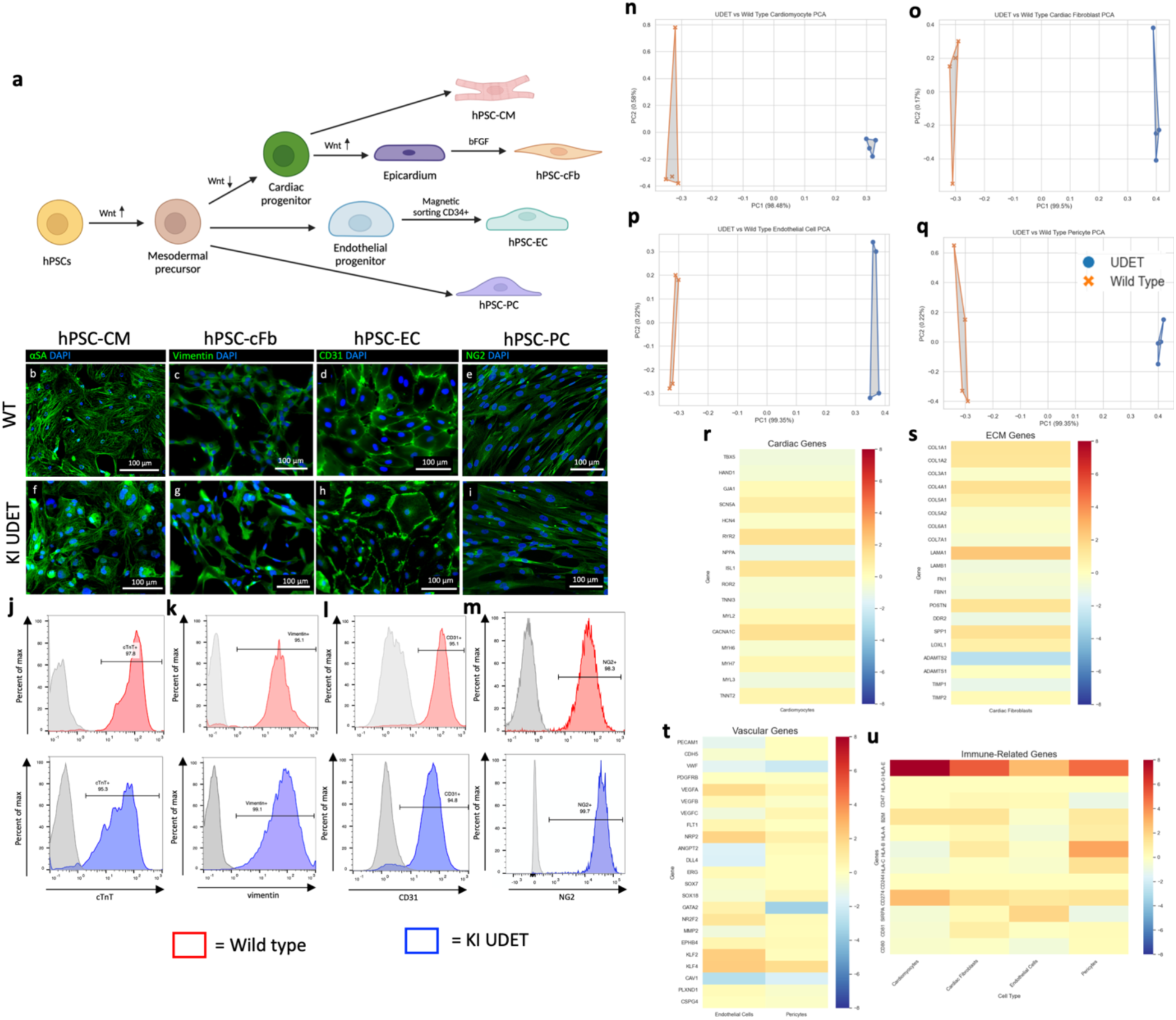
Differentiation of WT and KI UDET hPSCs into cardiac cell types. **a**, Schematic illustrating the differentiation of hPSCs into hPSC-CMs, hPSC-cFbs, hPSC-ECs, and hPSC-PCs. **b-i**, Immunofluorescent imaging of cell type-specific markers in (**b-e**) WT hPSC-derived cells and (**f-i**) KI UDET hPSC-derived cells with nuclei (blue) and (**b,f**) hPSC-CMs with αSA (green), (**c,g**) hPSC-cFbs with vimentin (green), (**d,h**) hPSC-ECs with CD31 (green), and (**e,i**) hPSC-PCs with NG2 (green). **j-m,** Flow cytometry of WT hPSC-derived cells (red) and KI UDET-derived cells (blue) showing (**j**) hPSC-CMs stained with cTnT, (**k**) hPSC-cFbs stained with vimentin, (**l**) hPSC-ECs stained with CD31, and (**m**) hPSC-PCs stained with NG2. Corresponding isotype controls shown in gray. **n-q**, PCA representing separation of WT and KI UDET (**n**) hPSC-CMs, (**o**) hPSC-cFbs, (**p**) hPSC-ECs, and (**q**) hPSC-PCs. **r-u,** Heat maps of differentially expressed genes between WT and KI UDET hPSC-derived cells related to (**r**) hPSC-CMs and cardiac genes, (**s**) hPSC-cFbs and ECM genes, (**t**) hPSC-ECs and hPSC-PCs and vascular genes, and (**u**) all four hPSC-derived cardiac cell types and immune related genes.

### KI UDET hPSC-derived cardiac cells evade immune cell-mediated lysis

Characterization of HLA expression in hPSC-derived cells confirmed the robustness of our hypoimmunogenic gene edits: the expression of HLA-A2 and HLA-B/C remained at <1% and the expression of HLA-E remained at over 95% for all KI UDET hPSC-derived cell types (**Fig. 4a-d and Supplementary** Fig. 9a-f). We then evaluated the capacity of the engineered HLA expression to induce immune evasion in assays measuring T cell- and NK cell-mediated lysis. CD8+ T cells were co-cultured with WT hPSC-derived embryoid body (EB) cells (**Fig. 4e**) to selectively expand alloreactive CD8^+^ T cells (**Supplementary** Fig. 10a), which were subsequently cultured with hPSC-derived cardiac cells (**Fig. 4e**). These allogeneic CD8^+^ T cells recognized and lysed KI UDET hPSC-derived cells to a significantly lower extent than their WT counterparts (**Fig. 4f-I and Supplementary** Fig. 10b-e). Similarly, when co-culturing NK and hPSC-derived cardiac cells (**Fig. 4j**), we observed KI UDET-derived cell populations were more resistant to NK cell-mediated lysis than their WT counterparts, with significantly less cytotoxicity observed in hPSC-CMs, hPSC-cFbs, and hPSC-PCs (**Fig. 4k-n, I and Supplementary** Fig. 10f-i). Interestingly, while lysis of KI UDET-derived hPSC-ECs trended lower, it was not significant. The killing of both WT and KI UDET hPSC-ECs (**Fig. 4m**) was overall higher than other cell types; therefore, we postulate that there are likely other stress-inducing ligands on hPSC-ECs promoting NK cell-mediated lysis^38^.

**Fig. 4:**
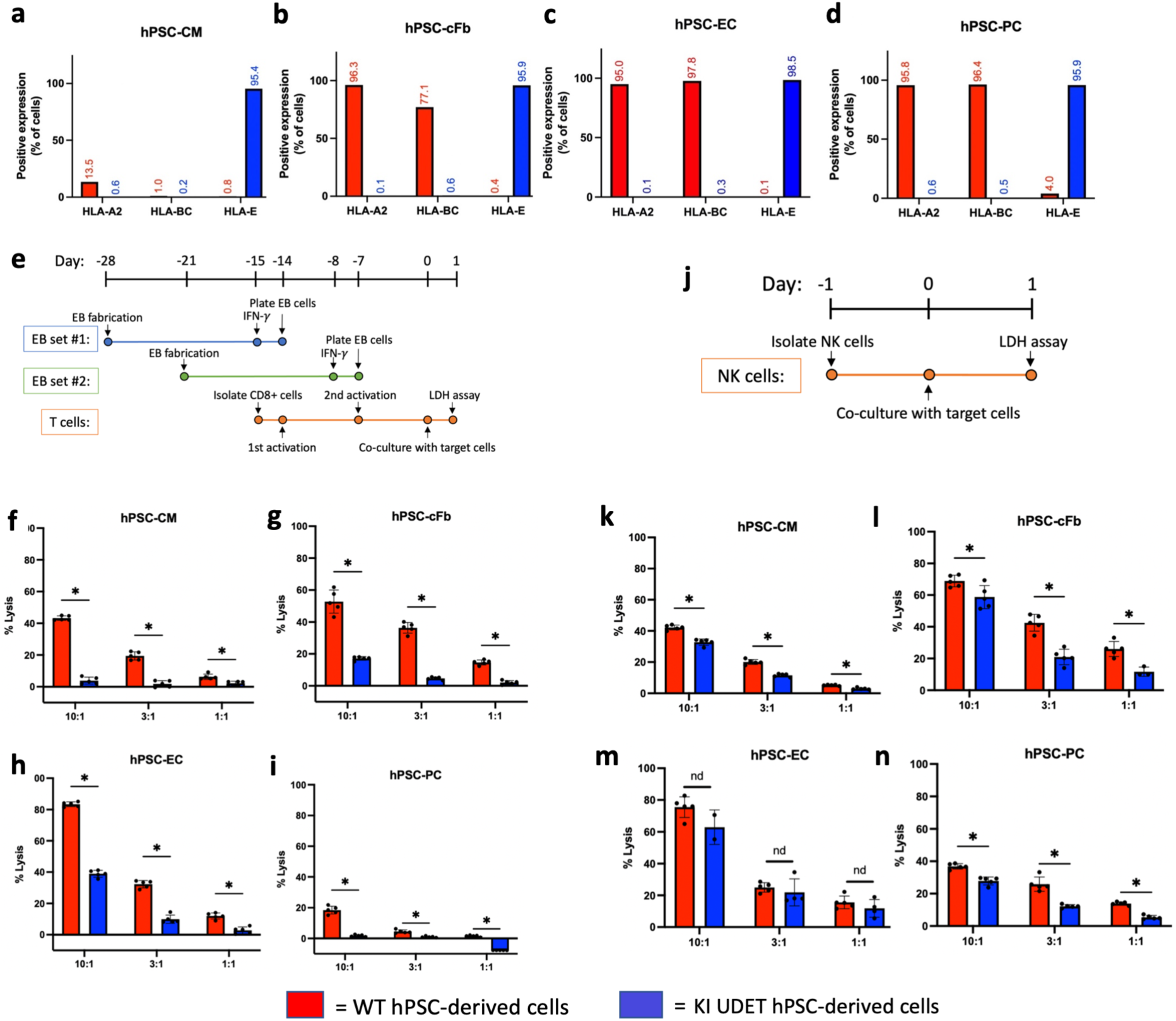
*In vitro* assessment of WT and KI UDET hPSC-derived immunogenicity. a-d, Bar charts of flow cytometry data showing expression of HLA-A2, HLA-BC, and HLA-E as percent positive expression in (**a**) hPSC-CMs, (**b**) hPSC-cFbs, (**c**) hPSC-ECs, and (**d**) hPSC-PCs by wild-type hPSC-derived cells (red) and UDET hPSC-derived cells (blue). **e**, Timeline of T cell cytotoxicity assay to measure T cell-mediated lysis of target cells. **f-i**, Percent lysis as quantified from LDH assay following exposure of activated CD8+ T cells to wild-type (red) and UDET (blue)-derived (**f**) hPSC-CMs, (**g**) hPSC-cFbs, (**h**) hPSC-ECs, and (**i**) hPSC-PCs. **j**, Timeline of NK cytotoxicity assay to measure NK cell-mediated lysis of target cells. **k-n**, Percent lysis as quantified from LDH assay following exposure of NK cells to wild-type (red) and UDET (blue)-derived (**k**) hPSC-CMs, (**l**) hPSC-cFbs, (**m**) hPSC-ECs, and (**n**) hPSC-PCs. Data shown from one donor representative of 2 biological replicates with 3-5 technical replicates. nd=no difference, *=p<0.05.

### Design and fabrication of KI UDET hPSC-derived cardiac organoids

While dissociated hPSC-CMs have demonstrated therapeutic potential, 3D cardiac microtissues have shown unique strength for implantation due to the enhanced cell-cell and cell-ECM interactions^61^. We therefore explored the fabrication of engineered cardiac organoids comprised of a defined population of hPSC-derived cardiac cells (**Fig. 5a**) based on existing literature demonstrating engineered tissue of this composition recapitulates hallmarks of cardiac tissue functions^62,63^. We confirmed that both WT and KI UDET cardiac cell populations self-aggregated to form 3D microtissues with similar spontaneous contractile capacity (**Fig. 5b**). We observed that both WT and KI UDET hPSC-derived cardiac organoids organized with hPSC-CMs in the peripheral region of the organoid and vimentin^+^ cells throughout the hPSC-CMs (**Fig. 5c,d**). Furthermore, hPSC-ECs formed lumenized CD31^+^ networks in the interior region of the organoids (**Fig. 5e,f**). PCA of bulk RNA-sequencing data demonstrated that PC1 accounted for 94% of the variance between WT and KI UDET hPSC-derived cardiac organoids, with HLA-E driving PC1 separation (**Supplementary** Fig. 11a,b), replicating our 2D cardiac cell findings (**Fig. 3n-q**). Additionally, DEA demonstrated subtle differences between WT and KI UDET hPSC-derived cardiac organoids with significantly higher expression of HLA-E in KI UDET hPSCs (**Supplementary** Fig. 11c,d). Total study PCA of hPSC-derived cardiac cells and organoids revealed that WT and KI UDET samples repeatedly cluster closely, with cardiac organoids positioned close to the cellular compositional average of 4 constituent cell types used for organoid fabrication (**Fig. 5g**). To investigate transcriptional differences influenced by the microenvironment provided in 3D cardiac organoids, we employed PCA to compare 3D organoids to 2D hPSC-CMs. This revealed that the addition of supporting cells drove separation between hPSC-CMs and cardiac organoids of both cell lines on PC1, with genes related to ECM development driving much of the variation (**Fig. 5h,i**). We observed distinct, paralleling separation on PC2 between WT and KI UDET samples (**Fig. 5h**), revealing HLA-E as the top gene loading of PC2 (**Fig. 5j**). Subsequent Gene ontology (GO) enrichment analysis of PC1 highlighted pathways associated with ECM organization and cell-substrate adhesion, supporting their critical roles in organoid formation (**Fig. 5k**). These findings prompted us to conduct DEA between 2D hPSC-CMs and hPSC-derived 3D cardiac organoids to highlight cardiac organoid-driven enhancement of key biological processes to promote graft survival and integration within the ischemic heart (**Fig. 5l**). We observed increases in genes related to vascular development and ECM production with similar patterns of enrichment between WT and KI UDET samples. To account for the cellular composition differences between 2D hPSC-CMs and 3D cardiac organoids, we generated a dataset comprised of transcriptomic data proportionally compiled from 4 constituent cell types (termed “*in silico* 2D hPSC cardiac cell aggregates”) (**Supplementary** Fig. 12a). PCA analyses demonstrated close clustering of *in* silico 2D aggregates to 3D cardiac organoids (**Supplementary** Fig. 12b,c) and revealed a strong enrichment of genes in 3D cardiac organoids compared to *in silico* 2D aggregates, with GO analysis indicating muscle tissue development to be strong drivers of gene enhancement of cardiac organoids (**Fig. 5m,n**). To assess cell-cell and cell-matrix interactions in 3D cardiac organoids, CellChat was used to interpret single cell RNAseq cardiac organoid datasets, revealing that hPSC-ECs and hPSC-PCs mediate cardiac development pathways via BMP and FGF signaling, while collagen-based cell-ECM interactions emphasize the critical roles of supporting cells aiding in graft retention (**Supplementary** Fig. 13 **and Fig. 5o,p**).

**Fig. 5:**
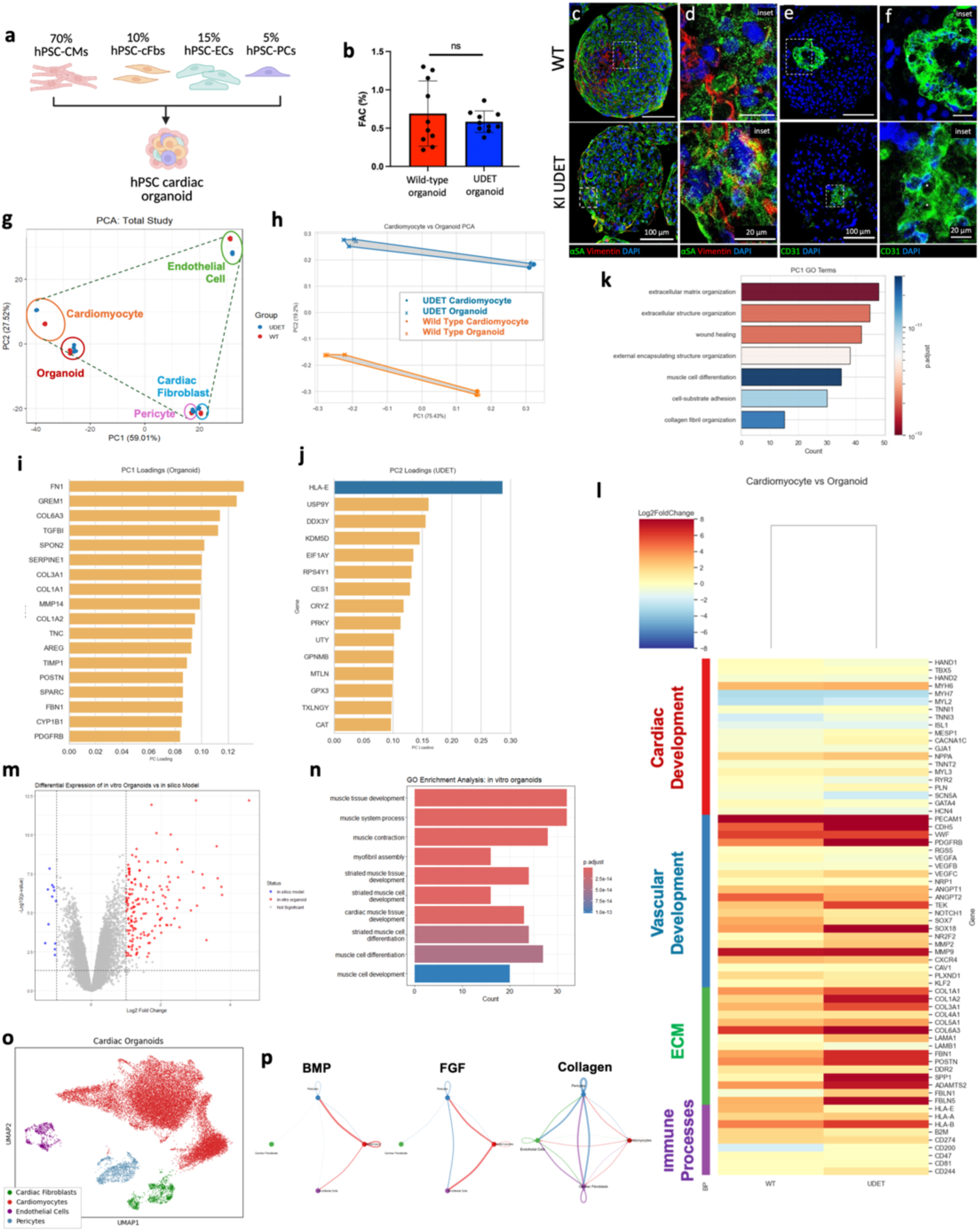
Development and characterization of WT and KI UDET hPSC-derived cardiac organoids. **a**, Schematic illustrating the composition and fabrication of hPSC-derived cardiac organoids. **b,** Contractile capacity of WT and KI UDET cardiac organoids, measured by percent FAC**. c-f,** Immunostaining of **(c,d**) hPSC-CM (αSA, green) and vimentin+ cell (red) organization as well as (**e,f**) hPSC-EC (CD31, green) organization and lumen formation (*) in WT (top) and KI UDET (bottom) hPSC-derived cardiac organoids**. g,** PCA detailing 2D hPSC-derived cell types and hPSC-derived cardiac organoids, outlining organoid position relative to each 2D cell type. **h,** PCA representing WT and KI UDET hPSC-CMs and hPSC-derived cardiac organoids. **i**, Negative PC2 loadings used in generation of PC plot in **h**. **j**, Positive PC1 loadings used in generation of PC plot in **h**. **k**, GO enrichment analysis highlighting pathways enriched in PC1 of the PC plot in **h**. **l**, Heat map of differential expressed genes between 2D hPSC-CMs and hPSC-derived cardiac organoids, highlighting genes involved in cardiac development, vascular development, ECM production, and immune processes. **m**, Differential gene expression between *in silico* 2D hPSC cardiac cell aggregates and 3D hPSC-derived cardiac organoids. **n**, GO terms upregulated in hPSC-derived cardiac organoids as compared to *in silico* 2D aggregates. **o,** Annotated UMAP highlighting clustering of each cell population within cardiac organoids. **p**, Circle plots generated via CellChat interpretation of single-cell RNA sequencing data.

### Functional engraftment of hPSC-derived cardiac organoids and attenuation of left ventricular remodeling after ischemia/reperfusion (I/R) injury

To evaluate the therapeutic potential of WT and KI UDET cardiac organoids, we injected organoids into an athymic rat model of ischemia/reperfusion (I/R) injury previously established by our lab^36^ (**Fig. 6a**). We observed successful engraftment of both WT and KI UDET-derived organoids at Day 28 and confirmed engraftment by immunostaining of slow-skeletal troponin I (TnI), a marker of hPSC-CMs ^64^ (**Fig. 6b,c**). We observed reduced left ventricular (LV) wall motion in control untreated I/R-injured hearts (control) (**Fig. 6d**) when compared to that of WT and KI UDET-derived organoid-treated groups (**Fig. 6e,f**). Quantification of fractional shortening (FS) revealed significant improvements at Day 7 and Day 28 for organoid injection groups compared to control (**Fig. 6g and Supplementary** Fig. 14a). Furthermore, WT and KI UDET cardiac organoids promoted significant recovery of FS that was lost to I/R injury by Days 7 and 28 (**Fig. 6h**), resulting in significantly greater increases in FS from 24 hours post-injury compared to control **(Fig. 6i)**. Analysis of left ventricular end diastolic diameter (LVEDD) revealed mostly insignificant differences among groups, in agreement with our previous report^36^. (**Fig. 6j-k and Supplementary** Fig. 14b). However, both WT and KI UDET organoids significantly recovered left ventricular end systolic diameter (LVESD) (**Fig. 6l-m and Supplementary** Fig. 14c), supporting enhanced contractility after organoid implantation.

**Fig. 6:**
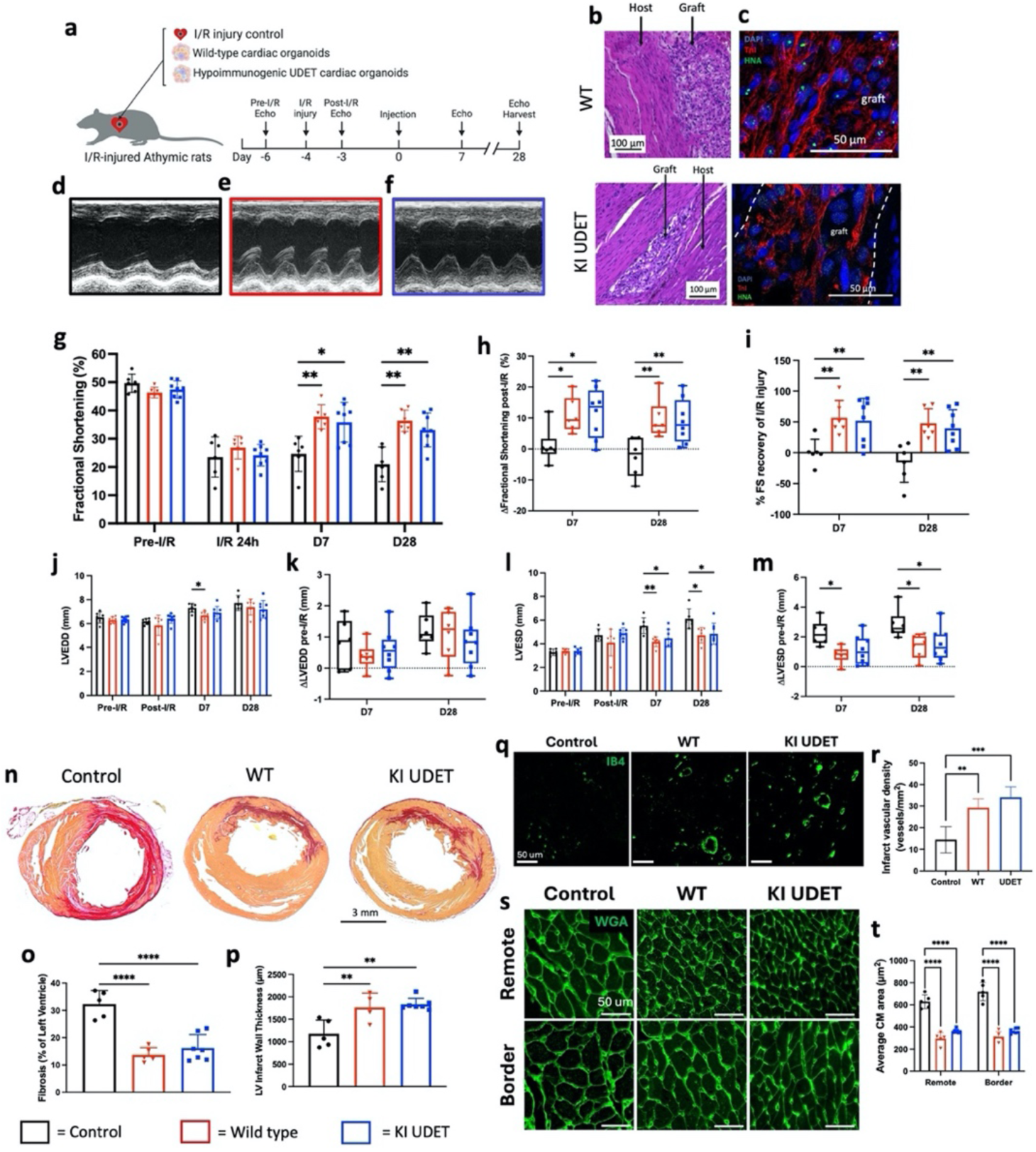
Engraftment, functional recovery, and attenuation of adverse left ventricular remodeling following injection of WT and KI UDET hPSC-derived cardiac organoids in I/R injured rat hearts a, Schematic showing injury control and organoid injections in I/R-injured athymic rats as well as experimental outline of organoid transplantation. **b**, H&E staining of engrafted organoids in WT (top) and KI UDET (bottom) organoid-treated rat hearts at Day 28. **c**, Immunofluorescent staining of slow-skeletal TnI (red) and human nucleolar antigen (HNA) (green) in WT (top) and KI UDET (bottom) organoid engraftments at Day 28. **d-f**, Representative M-mode echocardiographs at Day 28 from (**d**) control, (**e**) WT, and (**f**) KI UDET organoid-treated rat hearts. **g**, Average FS values at pre-I/R, post-I/R, and 7 and 28 days after transplantation. (n = 6, 6, and 8 rats for control, WT, and KI UDET organoid treatment groups, respectively). **h**, Average percent change in FS at Day 7 and Day 28 post-transplantation compared to post-I/R. **i**, Average percent recovery of FS lost to injury at Day 7 and Day 28. **j**, Average LVEDD values at pre-I/R, post-I/R, and 7 and 28 days after transplantation. **k,** Average change in LVEDD at days 7 and 28 compared to pre-I/ R. **l**, Average LVESD values at pre-I/R, post-I/R, and 7 and 28 days after transplantation. **m,** Average change in LVEDD at days 7 and 28 compared to pre-I/ R. **n**, Representative picrosirius red staining of infarcted rat hearts from control, WT organoid, and KI UDET organoid treatment groups (red, collagen I/III; yellow, myofibers). **o,** Average fibrosis of left ventricle at Day 28. **p**, Average LV wall thickness at Day 28. **q,** Vessels in the infarct zone at Day 28. **r**, Quantification of average infarct vascular density in hearts treated with primary cell or 4-cell isogenic organoids at Day 28. **s**, Representative images of wheat germ agglutinin (WGA)-stained host cardiomyocytes in border and remote zones of injured hearts at Day 28. **t**, Quantification of the average cross-sectional area of host cardiomyocytes in the border and remote zones of the host heart. *=p<0.05, **=p<0.01, ***=p<0.005, ****=p<0.001.

Hallmarks of pathological ventricular remodeling after myocardial infarction include cardiac fibrosis, LV wall thinning, and cardiomyocyte hypertrophy^65^. We identified collagenous scar tissue in all treatment groups, however, significant mitigation of fibrosis and left ventricular wall thinning was observed in WT and KI UDET treatment groups (**Fig. 6n-p**). Additionally, WT and KI UDET organoid transplantation correlated with significant increases in vascular density of infarcted myocardium (**Fig. 6q-r**). Finally, average cardiomyocyte area in remote and border regions of the left ventricle was significantly reduced in WT and KI UDET treatment groups (**Fig. 6s-t**), indicating less compensatory hypertrophy compared to control. Overall, our combined functional and morphological analyses demonstrate that hypoimmunogenic edits do not affect the therapeutic efficacy of KI UDET cardiac organoids.

### KI UDET hPSC-derived cardiac organoids evaded immune rejection in humanized mice

To assess the immunogenicity of KI UDET organoids *in vivo*, we transplanted hPSC-derived cardiac organoids under the kidney capsules of NSG mice previously humanized by human peripheral blood mononuclear cell (PBMC) engraftment (**Fig. 7a**). Immunostaining of cTnT indicated presence of both WT and KI UDET hPSC-derived cardiac organoids in grafted kidneys (**Fig. 7b**). While WT and KI UDET organoids were both surveilled by CD3^+^ T cells (**Fig. 7b**), we observed significant preservation of cTnT expression in KI UDET grafts (**Fig. 7c-d**). Furthermore, intragraft CD3 expression was significantly higher in WT organoid grafts compared to KI UDET (**Fig. 7e and Supplementary** Fig. 15), indicating that hypoimmunogenic cardiac organoids locally mitigate human T cell-mediated clearance *in vivo*. Finally, we determined systemic T cell responses to WT and KI UDET organoids and found that KI UDET transplantation correlated with fewer T cells in the blood (**Fig. 7f**) and significantly lower T cell expansion than WT when compared to their respective baselines (**Fig. 7g**). Interestingly, we observed a significant rise in CD8^+^ T cell frequencies and a decrease in CD4^+^ T cell frequencies in the blood of WT organoid-grafted mice 1 week after transplantation compared to those of KI UDET, which remained constant over time (**Supplementary** Fig. 16a,b). Taken together, our results demonstrate that KI UDET organoids effectively evade significant immume cell-mediated clearance and are well tolerated by a functional allogeneic human immune system.

**Fig. 7:**
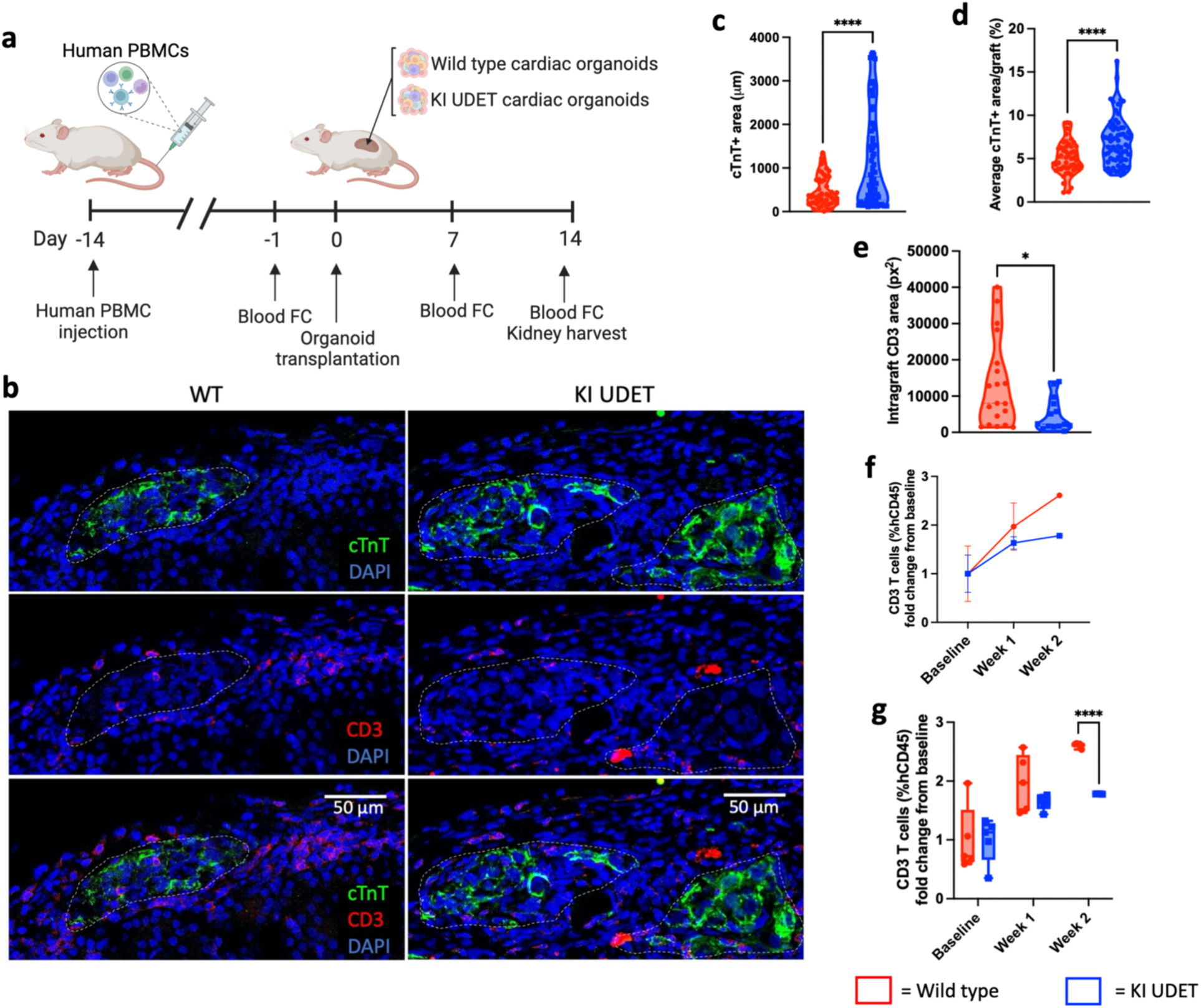
Kidney capsule transplantation of WT and KI UDET hPSC-derived cardiac organoids. **a**, Schematic illustrating timeline of humanization of NSG mice with human PBMCs as well as organoid transplantation and downstream analyses, FC = flow cytometry. **b**, Immunostaining of hPSC-CMs in cardiac organoid grafts (green, cTnT) and infiltrating T cells (red, CD3) in kidney capsules of mice injected with (left) WT cardiac organoids and (right) KI UDET hPSC-derived cardiac organoids. Dashed lines outlining graft area. **c-e**, Quantification of immunostaining displaying (**c**) cTnT+ area, (**d**) average cTnT+ area per graft, and (**e**) intragraft CD3+ area. **f-g**, Quantification of flow cytometry data evaluating CD3+ cells in circulation as percent of hCD45+ cells in mice receiving cardiac organoids kidney capsule transplantations, displayed as fold change from baseline. *=p<0.05, ****=p<0.001.

## DISCUSSION

Genetic mismatch between patients and stem cell-derived therapeutic cells is a major hurdle for the widespread implementation of regenerative medicine due to allogeneic immune rejection and the various toxic side effects associated with immunosuppressive drugs currently needed to stave off allogeneic rejection. By integrating CRISPR-Cas9 genome editing with engineered organoid technology, we provide the first demonstration of the therapeutic potential of hypoimmunogenic hPSC-CMs in the cardiac organoid platform for heart repair. To date, notable progress has been made in deriving of hypoimmunogenic islet cells to treat type 1 diabetes using HLA or B2M deletion and CD47 overexpression^40,44,45,66^. We chose HLA-E as an NK cell suppressor due to its suitability for cardiac applications. This was informed by the sizeable proportion of hPSC-CMs expressing the CD47 receptor (SIRPA)^67^, which could bind to CD47 and limit its ability to prevent NK cell-mediated lysis. In addition, our data highlight the importance of including presenting peptides in the trimer design to ensure robust, non-inducible expression of HLA-E and HLA-G on the cell surface. Using a single gene KO and single gene KI, our platform provides a straightforward and robust approach to derive stable universal donor hPSC lines for cardiac cell therapies. Our in-depth transcriptomic studies of 2D hPSC cardiac cells and 3D organoids revealed minimal off-target effects, supporting the precision of our genome engineering strategies and the lack of intracellular signaling of HLA-E^68,69^. These findings were further supported by structural and functional assessments, confirming that hypoimmunogenic editing preserved the integrity and therapeutic efficacy of hPSC derived cardiac cells and organoids.

Although organoids have been derived through guided differentiations of hPSCs^70–72^, our engineered organoid platform leverage the self-assembly of pre-differentiated hPSC-derived cardiac cells. This modular approach enables fine-tuning of organoid properties for specific applications, as we have previously demonstrated for disease modeling and cardiac cell therapy^36,73,74^. Here, we show that this platform is adaptable to include gene-edited cardiac cells that evade immune attack while maintaining their cardiac regenerative potential. Furthermore, our transcriptomic analyses underscore the functional benefit of incorporating each constituent cardiac cell type in a 3D organoid platform, revealing upregulations in pathways that may enhance cell survival after transplantation compared to dissociated 2D cell types. Taken together, our study integrates precise genome editing with engineered cardiac organoid technology to create hypoimmunogenic hPSC cardiac organoids. These findings establish a foundation for off-the-shelf regenerative cell therapies to treat cardiovascular diseases—the leading cause of death worldwide.

## METHODS

### Maintenance of hPSCs

Human pluripotent stem cells (H9, 6-9-9, 19-9-11 and IMR90) were maintained on either Matrigel (Corning) or iMatrix-511 silk (Iwai North America) coated plates in mTeSR1 (STEMCELL Technologies) medium according to previously published methods^75,76^. hPSCs with the GAPDH-Luc KI were drug selected until a stable KI line was established with 200 µg/mL geneticin. All drugs were removed upon initiating differentiation. Cells were routinely tested to ensure mycoplasma free culture conditions using an established PCR based detection method^77^. The use of hESCs was approved by the Institutional Review Board of the Pennsylvania State University and the Embryonic Stem Cell Research Oversight Committee. All studies were conducted in accordance with the approved guidelines.

### Maintenance of HEK293 cells

Human embryonic kidney (HEK) 293 cells were maintained in DMEM (Thermo Fisher Scientific) supplemented with 10% FBS (VWR) and passaged every three to four days via TrypLE Express (Thermo Fisher Scientific) mediated dissociation. Cells were routinely tested to ensure mycoplasma free culture conditions using an established PCR based detection method^77^.

### Maintenance of human peripheral blood-derived CD8^+^ T cells

These cells were maintained using ImmunoCult-XF T-cell Expansion Media (STEMCELL Technologies) and activated using ImmunoCult HuCD3/CD28/CD2 T-cell Activator (STEMCELL Technologies). The cells were cultured per the instructions provided by STEMCELL Technologies regarding the use of their media and activator. These cells were maintained for no longer than 21 days in culture.

### Maintenance of NK-92 cells

These cells were cultured in MEM-alpha supplemented with 0.2 mM myo-inositol (Sigma), 0.1 mM 2-mercaptoethanol (Gibco), 0.02 mM folic acid (Sigma), 100 IU/mL IL-2 (Peprotech), 12.5% FBS, 12.5% horse serum (Thermo Fisher Scientific). Fresh media was added every 2-3 days, and cells were gently pipetted to disperse clusters. Cell density was monitored to ensure it remained between 1x10^5^-1x10^6^ cells/mL at all times. Approximately every 5 days the cells were passaged by collection, centrifugation, and resuspension in fresh media. Cells were routinely tested to ensure mycoplasma free culture conditions using an established PCR based detection method^77^.

### Germ layer differentiation

For ectoderm differentiation: Cells were plated at low density (20% confluent). On day 0 and every day after, media was changed with LaSR Basal media for up to 6 days. For mesoderm differentiation: Cells were cultured until 80% confluent. On day 0, media was changed with RPMI (Thermo Fisher Scientific) supplemented with 100 µg/mL L-ascorbic acid and 6 µM CHIR99021. On day 1 media was changed to RPMI supplemented with 200 µg/mL L-ascorbic acid and 0.5% HSA (Biological Industries, 10% solution). Cells were collected on day 1 or 2. For endoderm differentiation: Cells were cultured until 60% confluent. On day 0, media was changed with RPMI supplemented with 1 µM dorsomorphin (Sigma) and 3 µM CHIR99021. Each day from day 1 to day 4, media was changed with RPMI supplemented with B-27 supplement (Thermo Fisher Scientific). Cells were collected on day 4.

### Generation of hPSC-CPCs and hPSC-CMs from IMR90 hPSCs

Cardiac differentiation of hPSCs was initiated when hPSCs seeded on Matrigel coated plates reached 80% confluence. Differentiation was performed according to previously published GiWi method^3,78,79^. Briefly, at day 0, cells were treated RPMI supplemented with 100 µg/mL L-ascorbic acid and 6 µM CHIR99021, followed by a change with RPMI supplemented with 200 µg/mL L-ascorbic acid and 0.5% HSA medium on day 1 and 2. On day 3, 2 µM Wnt-C59 (Tocris) was added, followed by a medium change with RPMI supplemented with 200 µg/mL L-ascorbic acid and 0.5% HSA on day 5. On day 6 and every 3 days after, cells were then cultured in RPMI supplemented with 200 µg/mL L-ascorbic acid, 0.25% HSA, selenium, transferrin, and insulin. Cells were considered cardiac progenitors between day 8 and day 12 and cardiomyocytes following day 12. Cells for *in vivo* transplantation were collected and counted on day 6.

### Generation of B2M KO sgRNA plasmid constructs

We designed two gRNAs, one targeted to exon 1 (CGCGAGCACAGCTAAGGCCA) and one targeted to exon 2 (TGTGAACCATGTGACTTTGTC). These gRNAs were ligated into a plasmid backbone, pGuide (This plasmid was a gift from Kiran Musunuru. Addgene #64711), that was linearized with BbsI (New England Biolabs). Sanger sequencing was used to confirm successful incorporation of the gRNA sequence. These plasmids are available as pLR05_pGuide_B2M_sgRNA1 and pLR06_pGuide_B2M_sgRNA2 from Addgene. The plasmid backbone for B2M sgRNA insertion was linearized using the BbsI (New England Biolabs) restriction enzyme. The linearization reaction was performed by incubating the reaction at 55°C (BsmBI) or 37°C (BbsI) for 5 hours and purifying the linearized plasmid from an agarose gel following electrophoresis using a Zymoclean Gel DNA Recovery Kit (Zymo Research, D4001).

The forward and reverse sgRNA DNA oligos (synthesized by IDT) were annealed and ligated into the linearized backbone. The ligated plasmid was transformed into Stbl3 competent E.coli (Thermo Fisher Scientific) and streaked on an Ampicillin containing agar plate (Thermo Fisher Scientific). A liquid culture was inoculated for 4-8 E.coli colonies and allowed to culture overnight. Plasmid was extracted from the E.coli using the Zyppy Plasmid Miniprep Kit (Zymo Research, D4020), and Sanger Sequencing was performed to identify a successfully ligated plasmid containing the sgRNA sequence. Sanger Sequencing was performed by the Penn State Huck Institutes Genomics Core Facility staff.

### Lipofection of HEK293 cells

Lipofectamine 3000 reagent (Thermo Fisher Scientific) and 3 µg total of the desired plasmid DNA were combined in basal media. This solution was then added to cells. Media was changed the next day, washing the cells once with culture media to remove the lipofectamine reagents. All plasmid DNA used was prepared using the Invitrogen PureLink HiPure Plasmid Filter Midiprep Kit.

### Generation of B2M KO cells

Cells were dissociated with Accutase for 10 minutes at 37C and pelleted. The cell pellet was resuspended in P3 Solution (Lonza) with 16 µg of plasmid DNA, including 7 µg CAG-Cas9-T2A-EGFP-ires-puro (This plasmid was a gift from Timo Otonkoski. Addgene #78311), 7 µg pLR05_pGuide_B2M_sgRNA1, and 2 µg pCE-mp53DD (This plasmid was a gift from Shinya Yamanaka. Addgene #41856). We used p53DD due to its ability to improve cell survival by inhibiting p53-mediated apoptosis and cell cycle arrest during Cas9 editing^57,58^, The mixture was transferred to a cuvette and nucleofected using the CB150 program on the Lonza 4D Nucleofector. All plasmid DNA used was prepared using an Invitrogen PureLink HiPure Plasmid Filter Midiprep Kit. Cells were plated at a high density with 5 µM Y27632. The next day media was changed and for 1-2 days following 1 µg/mL puromycin was added to the media to select for cells that were successfully nucleofected. Cells were then plated at a single cell density and single cell derived colonies were picked and expanded. Single cell derived colonies were tested via flow cytometry to identify successful homozygous KO clones. These clones were then further characterized, and allele sequencing was performed using PCR amplified DNA of the Cas9 cut site and the TOPO TA Cloning Kit (Thermo Fisher Scientific) following the manufacturer’s instructions.

### Generation of lentiviruses

To generate lentiviruses, the cargo plasmid and 2^nd^ generation packaging plasmids psPAX2 and pMD2.G were added into OptiMEM (Thermo Fisher Scientific) and incubated at room temperature for 5 minutes. psPAX2 and pMD2.G were gifts from Didier Trono (Addgene plasmid# 12260 and 12259). FuGENE HD reagent (Promega) was added to the mixture by careful pipetting and incubating at room temperature for an additional 10-15 minutes. All plasmid DNA used were prepared using an Invitrogen PureLink HiPure Plasmid Filter Midiprep Kit (Thermo Fisher Scientific). This solution was then added into pre-warmed DMEM+10% FBS to make the transfection media. Media was changed on 90% confluent HEK293 cells using 0.5X the normal culture volume of transfection media. Cells were incubated overnight at 37°C and media was changed the following morning with pre-warmed DMEM with 10% FBS using 1.5X the normal culture volume. 24hrs, 48hrs, and 72hrs after this media change, virus containing media was collected from the cells and the media was changed with pre-warmed DMEM with 10% FBS using 1.5X the normal culture volume. The cells were disposed of after the third collection.

The collected virus containing media was stored at 4°C until collection was complete. Virus containing media was centrifuged for 5 minutes at 1.5 G’s and the supernatant was collected. Concentration was done using Lenti-X Concentrator (TaKaRa Bio USA) according to the manufacturer’s instructions. Concentrated virus was resuspended in DMEM and stored in single use aliquots at -80°C until used.

### Lentiviral transduction of HEK293 cells

To transduce HEK293 cells, 200uL of fresh virus containing media is added directly into the culture media of those cells and gently pipetted to mix.

### Lentiviral transduction of hPSCs

To transduce hPSCs, an 80% confluent well of hPSCs was washed with DPBS and dissociated using Accutase for 5 minutes and pelleted. The supernatant was aspirated, and the cell pellet was resuspended in 50% culture media and 50% concentrated virus solution with a total volume of 100 µL. Cells were incubated at 37C for 30 minutes and then plated in prewarmed culture media with 5 µM Y27632 and incubated overnight. Media was changed 24h later.

### Generation of KI UDET and UDGT hPSCs

The dimer fusion proteins were designed by joining the B2M coding sequence (CDS) to the HLA-E or HLA-G CDS sequence by a flexible linker (GGGGS)x3. The trimer protein design was guided by previous research^80^. A nonamer peptide sequence for HLA binding and presentation, a flexible linker (GGGGS)x3, B2M, a flexible linker (GGGGS)x4, and the sequence for HLAE or HLAG was linked. For all fusion proteins, a point mutation (A267G) was made to the B2M sequence to remove a restriction site but maintain the amino acid sequence. The Kozak sequence was included at the start of each fusion protein. These fusion proteins were synthesized by Genewiz and cloned into lentiviral vector pXL001 (This plasmid was a gift from Sean Palecek. Addgene #26122) using restriction enzymes SpeI and EcoRI (New England Biolabs).

To generate the knockin donor plasmids, the trimer protein coding sequence was PCR amplified from pLR19_pXL001_CSIG-B2M-HLAE and pLR20_pXL001_PLASIG-B2M-HLAG using the Q5 High-Fidelity 2X Master Mix (New England Biolabs) and purified following gel electrophoresis. This sequence was ligated into AAVS1-Pur-CAG-EGFP (This plasmid was a gift from Su-Chun Zhang. Addgene #80945), which was linearized using restriction enzymes SalI and MluI (New England Biolabs), using the NEBuilder HiFi DNA Assembly Master Mix (New England Biolabs).

To generate the knockin cells, B2M KO cells were dissociated with Accutase (STEMCELL Technologies) for 10 minutes at 37C and pelleted. The cell pellet was resuspended in P3 Solution (Lonza) with 16 µg of plasmid DNA, including 7 µg eCas9-T2gRNA, 7 µg donor plasmid (either AAVS1_KIDonor_CSIG-B2M-HLAE or AAVS1_KIDonor_PLASIG-B2M-HLAG), and 2 µg pCE-mp53DD. The T2 sgRNA (GGGGGCCACTAGGGACAGGAT) was cloned into the eSpCas9(1.1) plasmid (This was a gift from Feng Zhang. Addgene #71814) which was linearized with restriction enzyme BbsI (New England Biolabs). All plasmid DNA used was prepared using an Invitrogen PureLink HiPure Plasmid Filter Midiprep Kit. The mixture was transferred to a cuvette and nucleofected using the CB150 program on the Lonza 4D Nucleofector. Cells were plated at a high density with 5 µM Y27632. Following puromycin selection to purify the successfully modified cells, single cell colony selection was performed, and PCR based genotyping was used to identify heterozygous knockin clones.

### Generation of IMR90 GAPDH-Luciferase Lines

The generation of our IMR90 GAPDH-Luciferase lines were conducted in line with the Kao *et al.* findings^81^. We inserted the luciferase downstream of the Glyceraldehyde 3-phosphate dehydrogenase housekeeping gene within our IMR90 universal donor cell lines (GAPDH-Luc)^81^ due to concern that the lentiviral luciferase reporter construct would get silenced after differentiation. Briefly, pGAPTrap-Luc2-IRESMeo, pGAPTrap-TALEN 1, and pGAPTrap-TALEN 2 (Addgene #’s 82509, 83368, and 83369, respectively) were purchased from Addgene. Plasmids were transformed into One Shot STBL3 E. Coli (Thermo Fisher Scientific), a single clone was picked, cultured, and plasmid DNA was purified at a high concentration using Invitrogen PureLink HiPure Plasmid Filter Midiprep Kit. When hPSCs reached ∼80% confluency, cells were collected and spun down. In the meantime, 100 µL P3 Solution (Lonza) was prepared in a 1.5 mL tube and mixed with 5 µg of pGAPTrap-TALEN 1, 5 µg pGAPTrap-TALEN 2, and 20 µg of pGAPTrap-Luc2-IRESMeo. Cell pellet was then resuspended in this plasmid DNA solution, transferred to a cuvette, and nucleofected using the CB150 program on the Lonza 4D Nucleofector. Cells were plated at a high density with 5 µM Y27632. Cells were then expanded and gradually drug selected to a final concentration of 200 µg/mL geneticin. Stable drug selected GAPDH-Luc cell lines were then verified using IVIS imaging. Two wells of these cells were changed with fresh mTeSR1 media immediately before imaging, either without or with 150 µg/mL D-luciferin (Promega). Luminosity was measured using the Caliper Life Sciences, IVIS Lumina LT Series III, and analyzed through the Living Image computer program.

### RNA sequencing

For hPSC-CM sequencing in **Supplementary** Fig. 1a, total RNA was isolated with Direct-Zol RNA Kits (Zymo Research, USA). RNA quality was checked using the BioAnlayzer 2100. Library preparation type used was Illumina TrueSeq Stranded mRNA, Poly-A selection. Samples were sequenced on an Illumina HiSeq2500. Briefly, the FASTQ sequence files were mapped to Human Genome, GRCh37 with STAR aligner. The outputted BAM alignment files were used for further analysis. The expression count table was produced by inputting the BAM files produced in a previous step and annotation file gencode.v27lift37 from Gencode into the FeatureCounts program.

For remaining bulk RNA-sequencing analyses, cells were matched for passage number between WT and KI UDET hypoimmune cell lines to reduce biological differences. Similarly, passage-matched WT and KI UDET hPSC-CMs, hPSC-cFbs, hPSC-ECs, and hPSC-PCs were used to fabricate WT and KI UDET cardiac organoids for RNA-seq analysis. Between 750,000 and 1 million cells were generated per replicate, per condition (n = 4) to ensure biological stability across conditions. To dissociate cardiac organoids gently, organoids were transferred into a conical tube, washed with PBS, and suspended in 2 mL TrypLE (Gibco) for 60 minutes at 37°C with mechanical stimulation at 5-minute intervals. Viability was verified via flow cytometry. Samples were sent to Novogene for sequencing and alignment to the hg38 reference genome.

BAM files were used to generate count matrices on the Galaxy Project online platform (v24.2.3) using featureCounts (v2.0.8). The RStudio (v2024.04.2) Raw count matrices were filtered for genes with less than 10 reads, then normalized using variance stabilizing transformation (VST) to account for size factor variation. The DESeq2 package (v.1.44.0) was used to perform differential expression analysis (DEA) between WT and KI UDET samples for each cell condition. DEA results for each cell condition were deemed significant with a log fold change > +/- 1.0 and an adjusted p-value of < 0.05. DEA results were filtered for key genes of various biological processes (cardiac development, vascular development, extracellular matrix function, and immune-related processes), and visualized using log fold change values via heatmap.

DEA was performed comparing cardiomyocytes (WT & KI UDET) versus cardiac organoids to ensure parallel enhancement of key genes across cell lines from cardiomyocyte to organoid culture. Following DEA, principal component analysis (PCA) was performed on VST-stabilized expression values for each cell condition to determine what constitutes differences between each cell type. PC loading scores of all genes were calculated to determine which genes drove variance between samples across PC1. The top 25 genes driving PC1-based variance between samples were visualized via bar plots. Gene Ontology (GO) enrichment analysis was performed across the top 500 genes driving variance toward organoid samples on PC1 (clusterProfiler, v4.12.6), with a p-value cutoff of 0.05 and q-value cutoff of 0.2 to filter for biologically relevant pathways.

### Flow cytometry analysis

Cells were prepared for flow cytometry via dissociation with 1X TrypLE Select for differentiated cells (Thermo Fisher Scientific) or Accutase for hPSCs. For intracellular stains, cells were centrifuged at 200×*g* for 5 min and resuspended in cold 1% Paraformaldehyde for fixation (20 min, 4°C). Cells were diluted 1:10 with 1X PBS and centrifuged at 200×*g* for 5 min. Cells were then rinsed once with Intracellular Flow Buffer (1X DPBS, 1% Bovine Serum Albumin (Thermo Fisher Scientific) and 0.1% Triton X-100 (Sigma-Aldrich)) and stored at 4°C until stained. Fixed cells were centrifuged (200×*g*, 5 min, room temperature) and resuspended in Intracellular Flow Buffer with primary antibodies. An unstained control group was incubated in only Intracellular Flow Buffer. Cells were incubated with primary antibody or Intracellular Flow Buffer overnight at 4°C. Post-incubation, all groups were rinsed with Intracellular Flow Buffer, centrifuged (200×*g*, 5 min, room temperature) and resuspended in Intracellular Flow Buffer with secondary antibodies and incubated for 1 h at room temperature, protected from light. Stained cells were rinsed twice with Flow Buffer and resuspended in 300 µL Flow Buffer for flow cytometry analysis. For cell surface markers, cells were centrifuged at 200×*g* for 5 min and resuspended in Flow Buffer (1X DPBS, 0.5% Bovine Serum Albumin, 2mM EDTA) with pre-conjugated antibodies and their corresponding isotype controls. Cells were incubated with antibody for 30 minutes at 4°C. Post-incubation, all groups with rinsed with Flow Buffer and centrifuged (200×*g*, 5 min, room temperature). Stained cells were resuspended in 300 µL Flow Buffer for flow cytometry analysis. Flow cytometry was performed using a MACSQuant Analyzer 10 (Miltenyi Biotec), Northern Lights Spectral Cytometer, or a BD Accuri C6 Plus flow cytometer, and analysis performed using FlowJo (FlowJo, Ashland, OR, USA). Primary antibodies used: OCT4 (1:100; Santa Cruz, sc-5279, clone C-10), NANOG (1:500; Thermo Fisher Scientific, PA1-097), SSEA4 (1:20; DSHB, MC-813-70), cTnT (1:200; Thermo Fisher Scientific, MA517192), vimentin (1:200; Abcam, ab8069, clone V9), CD31 (1:50; Thermo Scientific, MA529474), NG2, (1:200; Abcam, ab83178). Pre-conjugated antibodies used: B2M-APC (1:100; Biolegend, 316312, clone 2M2), HLA-A/B/C-PE (1:40; Biolegend, 311406, clone W6/32), HLA-E-APC (1:40; Biolegend, 342606, clone 3D12), HLA-G-APC (1:100; Biolegend, 335909, clone 87G), HLA-E-APC (1:50; Invitrogen, 17995342, clone 3D12HLA-E), HLA-BC-APC (1:50; Invitrogen, 17593542, clone B1.23.2), HLA-A2-PE-Cy7, clone BB7.2), CD25-APC (1:50; R&D Systems, FAB1020A, clone 24212), CD45-BV421 (1:50; Biolegend, 304032, clone HI30), CD3-BV605 (1:50; Biolegend, 344836, clone SK7), CD8-APC (1:50; Biolegend, 344722, clone SK1), CD4-PE (1:50; Biolegend, 980804, clone SK3). Secondary antibodies used: Goat anti-mouse Alexa 488 (1:1000; Thermo Fisher Scientific A-11029), Goat anti-rabbit Alexa 647 (1:1000; Thermo Fisher Scientific A-21244), Goat anti-mouse Alexa 647 (1:1000; Thermo Fisher Scientific A-21235), Goat anti-rabbit Alexa 647 (1:200; Abcam, ab150083), Goat anti-Mouse Alexa 647 (1:200; Abcam, ab150115), Goat anti-rabbit Alexa 488 (1:200; Abcam, ab150077), Goat anti-Mouse Alexa 488 (1:200; Abcam, ab150113).

### 2D fluorescent imaging

Cells were seeded and cultured under normal conditions in 24-well plates containing sterile glass coverslips. Once at an appropriate confluency for staining (70-100% confluent), wells containing coverslips were treated with 4% Paraformaldehyde (20 min, room temperature) for fixation. Wells were rinsed with 1X DPBS + Ca^2+^ + Mg^2+^ (Thermo Fisher Scientific) twice and stored in 1X DPBS + Ca^2+^ + Mg^2+^ at 4°C until ready for staining. All subsequent steps were performed on coverslips in the well plates. Antigen blocking was performed using 10% Goat Serum (Millipore Sigma) in PBST-0.01 (1X DPBS + Ca^2+^ + Mg^2+^ supplemented with 0.01% Triton-X) (Sigma-Aldrich) (1 h, room temperature) and then rinsed twice with PBST-0.01 containing Ca^2+^ and Mg^2+^. Cells were then stained with primary antibodies in PBST-0.01 for >16 h at 4°C: anti-B2M (1:100, Biolegend, 316302, clone 2M2), anti-PAX6 (1:20, DSHB), anti-T (1:100, R&D Systems, AF2085), anti-SSEA4 (1:100, DSHB, MC-813-70), OCT4 (1:100, Santa Cruz, sc-5279, clone C-10), NANOG (1:500, Thermo Fisher Scientific, PA1-097), anti-Vimentin (1:200; Abcam, ab8069, clone V9), anti-CD31 (1:50; Thermo Scientific, MA529474), anti-α-sarcomeric actinin (1:200; Abcloncal, a8939), anti-NG2 (1:200; Abcam, ab83178). Cells were rinsed twice with PBST-0.01 and stained with secondary antibodies for 1 h at room temperature, protected from light: Goat anti-mouse Alexa 488 (1:1000; Thermo Fisher Scientific A-11029), Goat anti-rabbit Alexa 647 (1:1000; Thermo Fisher Scientific A-21244), Goat anti-mouse Alexa 647 (1:1000; Thermo Fisher Scientific A-21235), Goat anti-rabbit Alexa 647 (1:200; Abcam, ab150083), Goat anti-Mouse Alexa 647 (1:200; Abcam, ab150115), Goat anti-rabbit Alexa 488 (1:200; Abcam, ab150077), Goat anti-Mouse Alexa 488 (1:200; Abcam, ab150113). Stained cells were rinsed once with PBST-0.01 and then stained with NucBlue nuclei stain (Thermo Fisher Scientific) diluted in PBST-0.01 according to the manufacturer’s instruction (20 min, room temperature, protected from light). Cells were rinsed twice with PBST-0.01 and then coverslips were removed from well plates. For mounting, coverslips were inverted onto microscope slides and mounted using Fluoroshield (Sigma-Aldrich). Images were taken using a BZ-X All-in-One Fluorescence Microscope (Keyence, Osaka, JP).

### Western blotting

Cells were washed with DPBS and lysed with Mammalian Protein Extraction Reagent (Thermo Fisher) with 1X Halt’s Protease and Phosphatase (Thermo Fisher) by incubation for 3 minutes. Cell lysate was collected and stored at -80 C until used. Samples were mixed with Laemmli sample buffer (BioRad) at a working concentration of 1X and incubated at 97 C for 5 minutes. Samples were loaded into a pre-cast MP TGX stain free gel (BioRad) and run at 200V for 30 minutes in 1X Tris/Glycine/SDS buffer (BioRad). Protein was transferred to a PVDF membrane using a Trans-blot Turbo Transfer System (BioRad). The membrane was blocked for 30 minutes at room temperature in 1X TBST with 5% Dry Milk. The membrane was incubated overnight at 4C with primary antibodies and for 1 hour at room temperature with secondary antibodies in 1X TBST with 5% Dry Milk: B2M (1:5000, 316302, Biolegend, clone 2M2). The membrane was washed between each antibody exposure with 1X TBST. Chemiluminescence was activated using Clarity Western ECL Substrate (BioRad) and the blot was imaged using a ChemiDoc Touch Imaging System and Image Lab software (BioRad). Blots were analyzed using Fiji software.

### *In Vitro* Luciferase release assay

Cardiac cells were counted and dispersed evenly into 96 well plates and allowed to adhere for at least 72 hours prior to initiating a lysis assay. At least one media change was done between cardiac cell plating and assay initiation. Just prior to the start of the assay, at least three wells dispersed across the plate were dissociated and counted to ensure even density was achieved. The cell counts for these wells were then averaged and to obtain a target cell number. Based on this, the number of effector cells required for 10:1, 5:1, and 2:1 effector to target ratios was determined. The luminescence was then measured to establish a baseline. Media was changed and 100 µL of effector cell media containing 150 µg/mL D-luciferin (Promega) was added. Cells were incubated for 5 minutes before luminescence was measured for the 0 hour time point. Effector cells were collected, pelleted and dispersed into wells (at least 3 wells for each ratio). Cells were then incubated together at 37C for 2hrs. After 2 hours, 150 µg/mL D-luciferin was added and cells were incubated for 5 minutes prior to obtaining the 2 hour luminescence measurement. Effector cells were resuspended in their normal culture media prior to dispersal for the assay. Cardiac cells in effector medium with no effector cells were used for a spontaneous death control. Cardiac cells in effector medium with 0.1% Triton-X (Sigma) were used as a positive control. All luminescence measurements were collected with a Tecan Infinite M Plex plate reader. Effector cells were collected, pelleted and dispersed into wells (at least 3 wells for each ratio). The percent of specific lysis was calculated using previously established methods^82^.

### *In vivo* humanized mouse study of hPSC-CPC immunogenicity

5-week-old male NOD.Cg-Prkdc^scid^ Il2rg^tm1Wjl^/SzJ (NSG; Jackson Labs, #005557) were purchased. Mouse husbandry was carried out in sterile conditions at Penn State University. All animal research was conducted in accordance with pre-approved Penn State University IACUC protocols.

Batches of day 6, CPCs from the IMR90 GAPDH-Luc WT, B2M KO, and HLA-ET KI lines were frozen in 5 x 10^6^ cell aliquots. For quality assurance, one well per batch were left to differentiate to > day 14 CMs, upon which Cardiac Troponin T (cTnT) flow was run for each batch to determine successful differentiation. Only batches with high percent cTnT positivity were used for subsequent transplantation.

6- to 8-week-old NSG mice were used for kidney capsule transplantation. The procedure for kidney capsule transplantation was carried out closely following previously reported methods^83,84^. Briefly, 5 million of the frozen CPCs were transplanted under the kidney capsule by thawing the cells, immediately loading into sterile PE-50 tubing, and delicately injecting into the kidney capsule. At least 1 month after transplantation, mice were i.p. injected with VivoGlo Luciferin (Promega, Cat. # P1042) at 150 mg Luciferin per Kg body weight and IVIS imaged using the Caliper Life Sciences, IVIS Lumina LT Series III. Mice that had a sufficiently strong signal indicative of a majority of the cells were successfully transplanted into the kidney capsule were then used for subsequent immune challenging, whereas mice that failed the surgery were euthanized.

Baseline day 0 bioluminescent IVIS images were taken of the mice again and 33.33 x 10^6^ human PBMCs (Stem Cell Technologies, Cat. #70025) were i.p. injected. PBMCs were thawed and immediately injected. One batch of 1 x 10^8^ PBMCs was equally divided between three mice, i.e., one mouse per condition, so that any variations between PBMC batches would be shared across each experimental condition. On days 7, 14, and 21 post PBMC i.p. injection, the mice were again IVIS imaged to plot the time course change in bioluminescent flux. Regions of interest were custom made for each mouse based on their day 0 IVIS images using the Living Image computer program. Using the same region of interest, day 0, 7, 14, and 21 fluxes were calculated. Relative change in flux to day 0 were then calculated for each mouse for each time point.

### hPSC-CM differentiation and culture

hPSC-CM differentiation (**Supplementary** Fig. 7a) was performed as described previously in^85^. The differentiation protocol was based on previously established methods, according to the GiWi protocol^79^ to establish cardiac progenitor cells. In brief, 90% confluent 19-9-11 hPSCs were singularized following incubation in Accutase for 6 minutes at 37°C, 5% CO_2_ and seeded onto Matrigel-coated 24-well plates at 600,000 cells/cm^2^ in mTeSR1 with 5 µM Y-27632 on Day -2. Media was exchanged with mTeSR1 24h later (Day -1). On Day 0, cells were treated with 6-8 µM CHIR99021 (Selleck Chemicals, Houston, TX, USA) in RPMI 1640 (Thermo Fisher Scientific) supplemented with 2% B27 Minus Insulin (Thermo Fisher Scientific, Waltham, MA) (RPMI/B27-). Precisely 24h later (Day 1), media was replaced with fresh RPMI/B27-. On Day 3, cells were treated with RPMI/B27-supplemented with IWP2 (Sigma-Aldrich). Precisely 48h later (Day 5), media was exchanged with fresh RPMI/B27-. By Day 6, cells were considered to be cardiac progenitor cells. On Day 7, media was replaced with RPMI 1640 supplemented with B27 with Insulin (Thermo Fisher Scientific) (RPMI/B27+). Media was subsequently replaced every other day. On Day 10, cells were purified via lactate purification^86^ for 48h. Lactate purification media consisted of glucose-free DMEM (Thermo Fisher Scientific) supplemented with 3mM Na-l-lactate (BeanTown Chemical, Hudson, NH, USA). On Day 12, media was replaced with RPMI/B27+ for 2 days of recovery from purification. On Day 14, purified hPSC-CMs were prepared for cryopreservation via dissociation using Accutase at 37°C, 5% CO_2_ for 40 minutes. Accutase was quenched 1:1 with RPMI/B27+ and centrifuged at 200×*g* for 5 minutes at room temperature. hPSC-CMs were resuspended in RPMI/B27+ supplemented with 30% Defined FBS (Cytiva Life Sciences, Marlborough, MA, USA), 10% DMSO (Sigma-Aldrich, St. Louis, MO, USA), and 5 µM Y-27632. hPSC-CMs were frozen overnight at -80°C in a Nalgene Mr. Frosty Freezing Container (Thermo Fisher Scientific) and transferred to liquid nitrogen for long-term storage. Thawing of cryopreserved hPSC-CMs consisted of seeding cells at 15,000 cells/cm^2^ on Matrigel-coated well plates in RPMI/B27+ supplemented with 10% Defined FBS and 5 µM Y-27632. Expansion of hPSC-CM^87,88^ began 24h after thaw via replacement of media with RPMI/B27+ supplemented with 3 µM CHIR99021. Media was exchanged with RPMI/B27+ supplemented with 3 µM CHIR99021 every other day for cardiomyocyte expansion.

### hPSC-cFb differentiation and culture

hPSC-cFb differentiation (**Supplementary** Fig. 7b) was performed as described previously in ^85^. In summary, 19-9-11 hPSCs were differentiated based on the GiWi protocol to derive cardiac progenitors^79^, followed by differentiation into hPSC-derived epicardial cells^89,90^ and finally into hPSC-cFbs. Briefly, Day 6 cardiac progenitor cells (see “hPSC-CM differentiation” above) were differentiated following seeding hPSCs on Day -2 at 550,000 cells/cm^2^ into Matrigel-coated 24-well plates and treated with 8-9 µM CHIR99021 on Day 0. For hPSC-derived epicardial cells, Day 6 cardiac progenitor cells were dissociated using Accutase at 37°C, 5% CO_2_ for 10 minutes and replated at 50,000 cells/cm^2^ in LaSR basal media, consisting of Advanced DMEM/F12 (Thermo Fisher Scientific), 10 µg/mL l-Ascorbic Acid (Sigma-Aldrich), 1X GlutaMAX (Thermo Fisher Scientific), and 5 µM Y-27632. On Day 7, cells were treated for 48h with 8 µM CHIR99021 in LaSR basal media. After 24h (Day 8), cells were treated with fresh LaSR basal with 8µM CHIR99021. After 48-h treatment, media was exchanged with LaSR basal media daily until Day 12. On Day 12, cells were considered to be differentiated into hPSC-epicardial cells and passaged 1:6 and/or cryopreserved in LaSR basal supplemented with 0.5 µM A83-01 (Abcam, Cambridge, UK), 10% FBS (Sigma-Aldrich), and 10% DMSO (Sigma-Aldrich). hPSC-epicardial cells were thawed in LaSR basal with 10% FBS, 0.5 μM A83-01 and 5 μM Y-27632 on Matrigel-coated plates. For maintenance of hPSC-epicardial cells, media (LaSR basal with 0.5 µM A83-01) was exchanged daily. For passaging, hPSC-epicardial cells were passaged at 1:6 once 90% confluent using TrypLE at 37°C, 5% CO_2_ for 10 minutes and replated in LaSR basal with 0.5 µM A83-01 and 5 µM Y-27632 onto Matrigel-coated 6-well plates. Cells were passaged 2-3 times before beginning hPSC-cFb differentiation. hPSC-cFb differentiation began once hPSC-epicardial cells reached 100% confluency on Matrigel-coated plates. The hPSC-epicardial cells were passaged 1:1 onto porcine heart extracellular matrix (HEM)-coated plates (Day −1). HEM-coated plates were prepared as described in ^85^. In brief, HEM solution at 10 mg/mL (preparation described in ^91^) was diluted in 0.02 M acetic acid to 20 µg/mL and added to well plates. Plates were incubated for >16 h at 37°C, 5% CO_2._ Prior to seeding cells, HEM solution was aspirated, and wells were rinsed with 1X DPBS (Thermo Fisher Scientific). hPSC-cFb differentiation began on Day 0 with 10 days of treatment with LaSR basal + 10 ng/mL βFGF (Invitrogen, Waltham, MA, USA). Media was changed daily. On Day 10, cells were considered to be hPSC-cFb and passaged onto HEM-coated well plates in FibroGRO Complete Media (Millipore Sigma, Burlington, MA, USA) with 2% FBS and 1X GlutaMAX. Cells were maintained in FibroGRO and passaged 1:6 using TrypLE when they reached 80–90% confluency.

### hPSC-EC differentiation and culture

hPSC-EC differentiation (**Supplementary** Fig. 7c) was performed as described previously in ^85^. hPSC-EC differentiation was based on previous reports^92–95^. Briefly, 80–90% confluent 19-9-11 hPSCs were dissociated with Accutase for 6 min at 37°C, 5% CO2, quenched with equal parts mTeSR1, and centrifuged at room temperature, 200×*g* for 5 minutes. Cells were resuspended in mTeSR1 with 5 μM Y-27632 and seeded at 125,000 cells/cm^2^ (Day -2). Media was replaced with mTeSR1 after 24 h (D-1). On Day 0, cells were treated with 6 μM CHIR99021 in LaSR basal for precisely 48 h, replacing with fresh LaSR basal with 6 µM CHIR99021 after 24 h. On Day 2, media was changed to LaSR basal and replaced daily. On Day 5, cells were dissociated with Accutase for 10 min at 37 °C, 5% CO2, quenched with LaSR Basal, and purified using CD34+ Microbeads (Miltenyi Biotec) following the manufacturer’s instructions. Purified CD34+ cells were seeded at 10,000/cm^2^ onto collagen I-coated wells (50 μg/mL) in EGM2 (PromoCell, Heidelberg, DEU) supplemented with 0.5 μM A83-01 and 10 μM Y-27632. Media was replaced 1 day after with EGM2 supplemented with 0.5 μM A83-01 (Day 6) and media was further exchanged every other day. On Day 10, cells were purified by treatment with Versene for 5 min to remove non-endothelial cells and rinsed with 1X DPBS. Cells were subsequently replated 1:3 onto fresh collagen I-coated plates after dissociation with TrypLE for 15 min at 37 °C, 5% CO2 and maintained in EGM2 supplemented with 0.5 μM A83-01.

### hPSC-PC differentiation and culture

hPSC-PC differentiation (**Supplementary** Fig. 7d) was based on the mesodermal pericyte induction protocol established by Faal et al.^96^. In brief, 90% confluent 19-9-11 hPSCs were singularized following incubation in Accutase for 6 minutes at 37°C, 5% CO_2_ and seeded onto Matrigel-coated 24-well plates at 200,000 cells/cm^2^ in mTeSR1 with 5 µM Y-27632 on Day -1. On Day 0, cells media was changed to Mesoderm Induction Media (MIM) (StemCell Technologies). MIM was replaced daily for 5 days. On Day 6, cells were replated onto 0.1% gelatin-coated plates in pericyte medium (ScienCell, Carlsbad, CA, USA). Pericyte media was replaced every other day until Day 10. On Day 10, cells were passaged at a concentration of 25,000 cells/cm^2^ and maintained in pericyte medium.

### Human NK cell and CD8^+^ T cell isolation

Human NK and CD8^+^ T cells were isolated from commercially available human peripheral blood mononuclear cells (PBMC) (STEMCELL Technologies) from HLA-A2 negative donors (as an allogeneic mismatch to WT hPSCs) using the NK (CD56^+^) Easy Sep Negative Isolation Kit (STEMCELL Technologies) and CD8^+^ Easy Sep Negative Isolation Kit (STEMCELL Technologies), respectively. Briefly, PBMCs were counted and resuspended in EasySep Buffer (STEMCELL Technologies) to a concentration of 5 x 10^6^ cells/mL in an acceptable volume range (0.5-2 mL). The isolation antibody cocktail was then added to the cells based on the volume of the cells (50 µL/mL) and mixed and incubated at room temperature for 5 min. Upon incubation, the magnetic Rapid spheres from the isolation kit were vortexed for 30 seconds and added to the sample at 50 µL/mL. The solution was topped up to 2.5 mL and mixed using a P1000. The sample was then placed into the EasySep Magnet (STEMCELL Technologies) and allowed to incubate for 3 min. After incubation, the EasySep magnet containing the tube with cells was inverted and poured into a sterile conical tube, yielding isolated human NK cells or CD8^+^ T cells.

### Cytotoxicity assay with human NK cells

Upon successful isolation of NK cells using the EasySep kit, the NK cells were cultured overnight in RPMI Complete Medium. Complete Medium consists of RPMI 1640, (Thermo Fisher Scientific), 10% FBS (Thermo Fisher Scientific), 1% GlutaMAX (Thermo Fisher Scientific), 1% Nonessential Amino Acids (Thermo Fisher Scientific), 1% 1M HEPES (Thermo Fisher Scientific), 1% Penicillin-streptomycin (Thermo Fisher Scientific), and 1% Sodium Pyruvate, and 20 ng/mL of IL-15 (Thermo Fisher Scientific). The following day, NK cells were counted and plated with either WT 19-9-11 or KI UDET cells that had been differentiated and plated at 10,000 cells/well in a 96-well flat bottom plate at n=3-5. Cells were plated at NK cell: Target cell (E:T) ratios of 10:1, 3:1, and 1:1 for 24 hours before performing CyQuant lactate dehydrogenase (LDH) cytotoxicity assays (Thermo Fisher Scientific)^97^.

### Embryoid body co-culture and cytotoxicity assay with human CD8^+^ T cells

On Day -28 of the experiment (**Fig. 4e**), an initial set of embryoid bodies (EBs) were formed from WT 19-9-11 hPSCs via a self-aggregation method in agarose molds and allowed to culture for 14 days before co-culture with T cells (Day -14). On Day -21, a second set of EBs were formed from 19-9-11 hPSCs via self-aggregation to prepare for the second round of T-cell activation. EBs were pretreated with 25 ng/mL of IFN-γ for 24 hours prior to the onset of co-culture with T cells (Day -15). On Day -14 of the experiment, the first set of EBs were collected and dissociated using TrypLE for 40 minutes at 37°C, 5% CO_2_, pipetting every 5 minutes. Dissociated cells from embryoid bodies were plated in a 48-well plate at 17,500 cells per well. Human CD8+ T cells (isolated on Day -15) were plated in the same well at a ratio of 4:1 (70,000 T cells/well). The coculture was maintained in KnockOut DMEM (Thermo Fisher Scientific) with 20% Fetal Bovine Serum (Thermo Fisher Scientific), 1%, Non-Essential Amino Acids (Thermo Fisher Scientific), 0.5% GlutaMAX (Thermo Fisher Scientific), and 0.1 mM of β-mercaptoethanol (Sigma-Aldrich). The culture was supplemented with 25 ng/mL of IFN-γ and 600 IU/mL of human Interleukin-2. Media was changed on the culture every other day, and the coculture was maintained for 7 days before the second round of stimulation with the second set of EB cells. On Day -7, the T cells were stimulated with another 17,500 EB cells that had been similarly pre-treated with IFN-γ for 24 hours (Day -8) and dissociated and plated in 48-well plates. On Day 0, the T cells were isolated from co-culture with EB cells by gently pipetting from the well. The T cells were then counted and plated against either WT or KI UDET cells that had been differentiated and plated at 10,000 cells/well in a 96-well flat bottom plate at n=3-5. Cells were plated at T cell: Target cell ratios of 10:1, 3:1, and 1:1 and incubated for 24 hours before performing CyQuant LDH cytotoxicity assays.

### Validation of antigen specific T cells via CFSE staining and flow cytometry

One well of T cells was loaded with carboxyfluorescien succinimidyl ester (CFSE) (Thermo Fisher Scientific), a cell proliferation tracker dye, following the manufacturer’s instructions. In brief T cells were resuspended in PBS, loaded with the CFSE, and allowed to incubate at 37°C for 20minutes in the dark. The cells were then quenched by adding 5 times volume of media containing 10% serum. The cells were then spun down and resuspended in pre-warmed media and allowed to incubate at 37°C for another 10 minutes to allow for hydrolyzation of the dye. T cells were then counted to account for CFSE induced toxicity and plated against EB cells as described in the previous section. After 4 days of culture, the cells were harvested by gently pipetting the cells in the well. The cells were centrifuged, resuspended, and washed in FACS Buffer (PBS with 5% FBS). The cells were then and assessed via flow cytometry on the Northern Lights Spectral Cytometer.

### CyQuant LDH cytotoxicity assay

At 24 h after the coculture between NK or T cells with target cells, 50 µl of each supernatant was harvested, and run through the CyQuant LDH Cytotoxicity Assay (Invitrogen) to determine the amount of cell lysis. Briefly, the supernatants were plated in a 96-well flat bottom plate, with target cells alone, and target cells treated with detergent as negative and positive controls, respectively. Substrate from the kit was added to the supernatants and allowed to incubate for 30 minutes in the dark. Stop solution from the kit was added, and then the plate was read at 490 nm and 680nm. Results were reported as percent lysis and were calculated using the following formula:

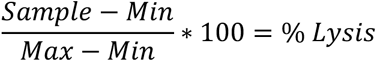

### hPSC cardiac organoid fabrication

hPSC cardiac organoids were fabricated based on previously published methods^36,85^. hPSC cardiac organoids were fabricated in a sequential seeding method (i.e., hPSC-CM spheroids (hPSC-CM only microtissues) were fabricated first, followed by seeding with supporting cell types to create hPSC cardiac organoids). To fabricate hPSC-CM spheroids, hPSC-CMs were expanded to super-confluency in RPMI/B27+ with 3µM CHIR99021 and allowed to recover in RPMI/B27+ for 24 h. hPSC-CM spheroids were fabricated on Day -14 (14 days prior to adding supporting cells). Agarose molds containing 35 microwells for organoid self-assembly were fabricated with 2% agarose, 24-h prior to seeding. Agarose molds were maintained in RMPI/B27+ to prevent drying. hPSC-CMs were dissociated using TrypLE for 40 minutes at 37°C, 5% CO_2_. hPSC-CMs were quenched and collected in RPMI/B27+, centrifuged (200×*g*, 5 min, room temperature), and resuspended to 4 x 10^6^ cells/mL in RPMI/B27+. For organoids for *in vitro* studies, 147,000 hPSC-CMs were plated into agarose molds (4,200 hPSC-CMs/microwell). For injection organoids, 70,000 hPSC-CMs were plated into agarose molds (2,000 hPSC-CMs/microwell). hPSC-CMs were permitted to settle in microwells for 20 min before adding RPMI/B27+ to each well. After 14 days of hPSC-CM spheroid culture with media changes every other day, supporting cells were added (Day 0). All cell types were dissociated using TrypLE (37°C, 5% CO2): 30 min for hPSC-ECs, 15 min for hPSC-cFbs, and 15 min for hPSC-PCs. Cells were quenched and collected in their cell-type specific maintenance media, centrifuged (200×*g*, 5 min, room temperature), and resuspended to 4 × 10^6^ cells/mL in organoid media. Organoid media consisted of 48% DMEM/F12 (Thermo Fisher Scientific), 5.4% Defined FBS, 0.54% Non-essential amino acids (Thermo Fisher Scientific), 30% Fibroblast Growth Media-3 (PromoCell), and 16% EGM2. Media was carefully aspirated from spheroid wells and supporting cell mixtures were prepared to plate onto hPSC-CM spheroids with the final cell ratio: 70% hPSC-CM: 10% hPSC-cFb: 15% hPSC-EC: 5% hPSC-PC. Cells were permitted to settle into the microwells of the agarose molds for 20 min then filled with organoid media. Organoids were maintained at 37°C, 5% CO2, and media was exchanged every other day.

### Video analysis of beating organoids

As described in previous reports^36,73,85^, videos of spontaneously beating organoids were recorded using a Carl Zeiss Axiovert A1 Inverted Microscope and Zen 2011 software (Zeiss) with a frame rate of 14 frames per second. To measure changes in area during contraction and relaxation, videos were converted to a sequence of TIFF format pictures with ImageJ (National Institute of Health). A threshold was applied to high contrast videos of contracting organoids and the area of the organoids was subsequently measured for each frame of the video. The changes in area during contraction and relaxation were measured as a percent change in fractional area change amplitude.

### Immunofluorescent staining of *in vitro* organoids

Organoid immunofluorescence staining was performed as previously described^36,85^. In brief, one well of organoids was embedded by flash-freezing in Optimal Cutting Temperature Compound (OCT) (Thermo Fisher Scientific) in Tissue-Tek plastic cryomolds (Sakura Finetek, Torrance, CA, USA), and immediately frozen to -80°C. Embedded blocks were sectioned at 8 µm per section using a Cryostat Microtome (Leica Biosystems, Wetzlar, DEU) and mounted on glass slides (Fisher Scientific, Pittsburgh, PA, USA). Mounted sections were stored at -20°C. Sections were then fixed with 100% cold Acetone (12 min) and allowed to dry. Sections were blocked with 10% Goat Serum in 1X DPBS with 0.01% Triton X-100 (PBST-0.01) (1 h, room temperature). Sections were then stained with primary antibodies diluted in PBST-0.01 for >16h at 4°C. Sections were then rinsed for 5 min with PBST-0.01 and then stained with secondary antibodies for 1 h at room temperature, protected from light. Sections were subsequently rinsed for 5 min with PBST-0.01 and then stained with NucBlue for 20 min at room temperature, protected from light. Sections were then rinsed twice for 5 min with PBST-0.01, and a coverslip was mounted using Fluoroshield. Primary antibodies used: anti-Vimentin (1:200; Abcam, ab8069, clone V9), anti-CD31 (1:50; Thermo Scientific, MA529474), anti-α-sarcomeric actinin (1:200; Abcloncal, a8939), Secondary antibodies used: Goat anti-rabbit Alexa 647 (1:200; Abcam, ab150083), Goat anti-Mouse Alexa 647 (1:200; Abcam, ab150115), Goat anti-rabbit Alexa 488 (1:200; Abcam, ab150077), Goat anti-Mouse Alexa 488 (1:200; Abcam, ab150113).

### *In silico* 2D hPSC cardiac cell aggregate development

An *in silico* model comprised of proportional aggregate data from each 2D cell type in hPSC-derived cardiac organoids was developed using relevant weighted bulk RNA-seq data of 2D cardiac cells (70% hPSC-CMs, 15% hPSC-ECs, 10% hPSC-cFbs, 5% hPSC-PCs). To develop these, we used vst-normalized gene reads across both WT and KI UDET lines for, averaging expression for each condition across biological replicates, then providing a weighted score by cell type to sum into the *in silico* gene expression profile for each gene. We then compared our silico data against bulk RNA-seq data of *in vitro* organoids via PCA, then performed DEA using the limma package (v.3.60.6) due to the normalized nature of the input data, which utilizes a linear model to perform differential expression. Significant differences via DEA were visualized via volcano plot, with all significant genes (162) for *in vitro* organoids used for GO Term Enrichment analysis.

### Single cell RNA-sequencing

We performed scRNA-seq on cardiac organoids at day 5 of culture. Libraries were prepared according to the 10X Genomics protocol, and libraries were sent to Azenta Life Sciences for sequencing. Reads were aligned to the human genome GRCh38. Default parameters were used to align reads and count unique molecular identifiers (UMIs) in order to create gene-cell expression matrices. The Python package Scanpy (version 1.10.2) was used to perform downstream scRNA-seq analysis. Datasets were filtered for low-quality cells, removing any cells with greater than 25% mitochondrial reads, less than 200 genes, and less than 3 cells, with any remaining outliers removed. Datasets were normalized, scaled, and visualized using Uniform Manifold Approximation and Projection (UMAP). Canonical markers for cardiomyocytes, cardiac fibroblasts, endothelial cells, and pericytes were used to annotate clusters (TNNT2, COL1A1, PECAM1, PDGFRB/RGS5). Once annotated, anndata (.h5ad) objects were saved for future use of downstream analyses such as CellChat. We used CellChat (version 2.1.2), an R package to analyze cell-cell interactions for canonical cell signaling pathways, to identify which cell types within our isogenic organoid were implicated in pathways of interest (https://github.com/sqjin/CellChat). Ligand-receptor networks and data visualization was carried out according to the vignette provided by CellChat.

### Rat model of ischemia-reperfusion injury

Procedures for our rat model of ischemia-reperfusion injury were similar to those previously described in ^36^. However, in this study, we used ketamine and xylazine for anesthesia instead of isoflurane to reduce post-operation mortality. Briefly, male athymic rats (8 to 10 weeks old, RNU Nude rats, Charles River) were anesthetized with ketamine and xylazine, intubated, and mechanically ventilated (SomnoSuite Low-Flow Anesthesia System, Kent Scientific). Rodent Surgical Monitor (INDUS instruments) was used to monitor the animal’s condition throughout the surgery, including heart rate, body temperature, and heart function through ECG. A left thoracotomy was performed to expose the heart. The pericardium was opened, and the left anterior descending coronary artery was ligated below the left atrial appendage level using a 7-0 Prolene suture (Ethicon). Occlusion was confirmed by LV blanching and ST elevation on ECG after ligation. After 1 h, the ligation was released, the chest cavity was aseptically closed, and the rat was allowed to recover for 24 h for echocardiographic measurement. For Day 28 studies, rats that met the echocardiographic inclusion criterion (FS <30%) were selected for experiments and randomly assigned into experimental groups.

### Injection of hPSC-derived cardiac organoids into injured rat hearts

Procedures for hPSC organoid injection were based on those described in ^36^. In brief, we injected organoids 4 days after I/R induction to allow angiogenesis-associated wound healing processes to reach their peak^10^. The rats that met our inclusion criterion were anesthetized with ketamine and xylazine, intubated, and mechanically ventilated. A second thoracotomy was performed to expose the heart. In preparation for injection, one-third of the total number of cardiac organoids was manually loaded into each of the three syringes with a Geltrex (Thermo Fisher Scientific) plug, which was used to occupy the dead space in the syringe for organoid injection and plug the injection path to prevent washout of organoids. A suspension of either WT or KI UDET hPSC cardiac organoids containing 500 organoids per rat for a total number of ∼1 x 10^6^ total hPSC-CMs and ∼1.5 x 10^6^ total cells per rat and were injected with a modified 29-gauge needle into the region surrounding the injured area at three sites (50 μL per site). I/R-injured rats without the cell injection were used as controls and did not receive a second surgery. Once the injections were completed, the chest was closed in three layers, and the animal was recovered with buprenorphine administered intraperitoneally to provide pain relief.

### Echocardiography

Echocardiography was performed as described in ^36^. Briefly, transthoracic echocardiography was performed using Vevo2100 ultrasound imaging system (VisualSonics, Toronto, Canada) with a 13 to 24 MHz linear array transducer (MS250). Rats were placed in supine position on a warming platform at 37°C under light (1 to 2%) isoflurane anesthesia to maintain the heart rate between 350 and 450 beats/min during imaging procedure. Cardiac function was examined using parasternal long-axis and short-axis 2D B-mode images and M-mode images at parasternal short-axis mid-papillary level. Offline image analyses were performed by an experienced investigator blind to the experimental groups using the dedicated Vevo LAB software. FS was calculated by the equation: FS = (LV end-diastolic dimension − LV end-systolic dimension)/LV end-diastolic dimension.

### Histological evaluation of harvested rat hearts

The following procedures were performed as described in ^36^. In brief, excised rat hearts were fixed overnight in 4% paraformaldehyde solution (Sigma Aldrich) and then to 70% ethanol before paraffin embedding. Using an automated microtome (LeicaRM2255, Leica, Exton, PA), paraffin-embedded hearts were sectioned at 8 μm in thickness. Slides were deparaffinized in xylene for 5 min and rehydrated using an ethanol gradient. Slides were then stained with hematoxylin and eosin (H&E) and picrosirius red stain kit (Abcam) or processed with heat-activated antigen retrieval solution (Vector Laboratories, Burlingame, CA) for 5 minutes for immunostaining as detailed below. H&E staining was performed every 100 µm to identify organoid grafts. Sections for immunostaining (described below) were selected as adjacent slides to H&E-identified organoid grafts. For percent fibrosis and wall thickness calculations, three regions from base to apex in each heart were used to quantify averages.

### Immunostaining of rat hearts

Immunostaining was performed as described previously^36^. Briefly, following heat-activated antigen retrieval, fixed rat heart sections were washed with PBS, permeabilized with 0.1% Triton X-100 (Sigma-Aldrich) in PBS (PBST-0.1) and blocked with 10% Goat serum (Sigma-Aldrich) for 2 h at room temperature. Primary antibodies were applied overnight, and secondary antibodies were applied for 1 h at room temperature, with 3, 10-min PBST-0.1 washes between primary and secondary stains. Nuclei were counterstained with DAPI (Molecular Probes/Invitrogen, Eugene, OR) diluted in PBST for 30 min at room temperature. Following the final wash procedure (PBST-0.1, three times for 5 min), glass coverslips were added to the slides using Fluoroshield. Lastly, a TCS SP5 AOBS laser scanning confocal microscope (Leica Microsystems Inc., Exton, PA) was used to acquire fluorescent images. Primary antibodies used: anti-TnI (1:200; Invitrogen, 710580), anti-HNA (1:200; Abcam, ab190710). Secondary antibodies used: Goat anti-rabbit Alexa 647 (1:200; Abcam, ab150083), Goat anti-Mouse Alexa 647 (1:200; Abcam, ab150115), Goat anti-rabbit Alexa 488 (1:200; Abcam, ab150077), Goat anti-Mouse Alexa 488 (1:200; Abcam, ab150113).

### Evaluation of cardiac hypertrophy and infarct vascular density

Cardiac hypertrophy measurements were performed as described previously^36^. In brief, cardiomyocyte cross-sectional area was determined via staining of cell boundaries with Alexa Fluor 488-conjugated WGA (Invitrogen). Slides were incubated with WGA (10 μg/ml) for 20 min at room temperature and washed with PBS three times before mounting coverslips. Three regions from base to apex in each heart were used to quantify average cross-sectional area per heart. To quantify the cell size, three independent samples per group with three different regions of interest from the border and remote zones in the left ventricle were captured at 40x magnification with a TCS SP5 AOBS laser scanning confocal microscope (Leica Microsystems). Fluorescent images were then processed by thresholding and individual cell areas were segmented and calculated using ImageJ batch processing. To quantify infarct vascular density, three independent sections from infarcted hearts in each group were stained with Isolectin GS-IB4, Alexa Fluor™ 568 Conjugate (Invitrogen). Lumenized IB4-positive vessels were counted manually in each image and converted to vessels/mm^2^.

### hPSC cardiac organoid transplantation under the kidney capsule in humanized mice

Human PBMCs were freshly isolated and 10 x 10^6^ PBMCs were retro-orbitally intravenously injected into each NOD-*scid*IL2Rg^null^ (NSG) mouse. After 14 days, confirmation of PBMC engraftment and sufficient mouse humanization was confirmed via bleeding and assessment of circulating hCD45+ cells via flow cytometry . In mice with confirmed humanization, WT and KI UDET hPSC-derived cardiac organoids were transplanted under the kidney capsule of humanized mice as previously described^98^ and in accordance to the IACUC of the Medical University of South Carolina (MUSC). In brief, mice were anesthetized with isoflurane followed by lubricant application on the eyes and subcutaneous injections of analgesics (buprenorphine and meloxicam). After shaving and applying Nair, the surgical area was sterilized with povidone-iodine and alcohol swabs. The surgery was then performed aseptically, first by making an incision in the skin and then in the peritoneal layer. The kidney was then exposed and approximately 200 organoids were transplanted into the kidney capsule for a total of 0.6 x 10^6^ cells per mouse. The kidney was returned to its original position, the peritoneal incision closed with surgical stitches, and the skin incision closed with surgical clips. Experimental animals were monitored and received additional subcutaneous injections of analgesics in the following days.

### Histological analysis of cardiac organoid graft-bearing mouse kidneys

Kidneys containing hPSC-derived cardiac organoids were processed as described in ^98^. In brief, kidneys were fixed in 4% PFA at room temperature for 2 h. The graft-containing kidney was then cut in half perpendicular to the graft without touching the graft and incubated in 7 mL 4% PFA for an additional hour at 4°C. The sliced fixed graft-containing kidney was then transferred to 5 mL 30% sucrose solution and incubated overnight at 4°C. The following day, the kidney halves were mounted in OCT medium in a cryomold and frozen on top of dry ice. OCT blocks were stored at −80°C until being sectioned and stained with antibodies. To assess organoid engraftment and T cell infiltration within cardiac grafts in a humanized mouse kidney capsule model, immunofluorescence images were analyzed using a custom Python-based pipeline. The green fluorescence channel (cTnT) was used to identify the graft region, while the red channel (CD3) indicated T cell presence. Evaluation of cardiac organoid engraftment via cTnT+ area was chosen because cTnT is a protein unique to our hPSC-derived cardiac organoids (i.e., in hPSC-CMs) not present in the kidney capsule, as well as the fact that hPSC-CMs make up the largest portion of our hPSC-derived cardiac organoids (70% hPSC-CMs). The cTnT signal was enhanced and binarized using a fixed intensity threshold, followed by morphological closing to fill gaps and removal of small objects to reduce noise. The red CD3 channel was similarly thresholded to identify regions of positive T cell signal. Intra-graft CD3 presence was defined as the overlap between the CD3+ mask and the refined graft mask. The intra-graft CD3 area was quantified by calculating the total number of CD3+ pixels located within the graft region. Primary antibodies used: anti-cTnT (1:200; Abcam, ab8295), anti-CD3 (1:200; Thermo Scientific, MA547763). Secondary antibodies used: Goat anti-rabbit Alexa 647 (1:200; Abcam, ab150083), Goat anti-Mouse Alexa 647 (1:200; Abcam, ab150115), Goat anti-rabbit Alexa 488 (1:200; Abcam, ab150077), Goat anti-Mouse Alexa 488 (1:200; Abcam, ab150113).

### Statistics

Data obtained from multiple experiments or replicates are shown as the mean ± standard error of the mean. Where appropriate, Student’s *t* test, one-way ANOVA, or a two-way ANOVA were utilized (alpha = 0.05) with a Bonferroni or Tukey’s post hoc test. Data were considered significant when p<.05. Statistical tests were performed using GraphPad Prism.

## ACKNOWLEDGEMENTS

This work was supported by the National Institutes of Health (NIH) (1R01HL133308, 1R01HL175050, 1U01HL169361, 1R01HL168255, 1R21HL167211, and 1R01HL173532), the National Science Foundation (NSF) Award #OIA-2242812, and American Heart Association Grant # 23IPA1054426/Ying Mei/2023 to Y.M.; NSF CBET-1943696 and NIH grants (R21EB026035, R56DK133147) to X.L.L.; NSF Center for Cell Manufacturing Technologies (CMaT; grant EEC-1648035), NSF grant CBET-2225300, and NIH grant R01HL165726 to S.P.P. This work was also supported by F31HL154665 to C.M. K., F31HL156541 to R.W.B., and F30HL160055 to D.C.A. We would like to thank Dr. Peters at Penn State University for access to his IVIS facility. This study used the services of the Morphology, Imaging, and Instrumentation Core, which is supported by NIH-NIGMSP30GM103342 to the South Carolina COBRE for Developmentally Based Cardiovascular Diseases and the Flow Cytometry and Cell Sorting Shared Resource at the Hollings Cancer Center, which is supported by NIH-NCIP30CA138313.

## AUTHOR CONTRIBUTIONS

S.E.S., A.R.H., D.C.A., L.N.R., X.L.L., and Y.M. conceived study details with assistance from all contributing authors. J.M.R. contributed to the study conceptualization and its clinical relevance. S.E.S., A.R.H., L.R., X.L.L. and Y.M. prepared the manuscript with assistance from all contributing authors. A.R.H. and L.N.R. performed hypoimmunogenic gene edit studies and wrote the manuscript. X.L.L. designed the project and performed the experiments. X.B. contributed to the design of gene editing experiment. hPSC cardiac differentiations were performed by S.E.S. and C.M.K. with assistance from M.E.F. and S.P.P. Organoid fabrications and characterizations were conducted by S.E.S. with assistance from C.M.K. N.A.H. performed bulk and single-cell RNA sequencing analysis, with sample preparation by S.E.S., M.L., J.E.M., and R.A.N. Immunological assays for cardiac cell types were performed by D.C.A., N.A.H. and S.E.S, with guidance from L.M.R.F. M.L. performed rat surgeries and echocardiographic studies. A.R.H. and L.M.R.F. performed mouse surgeries with assistance from R.A.R. and S.E.S. R.W.B., S.E.S., and J.D.B. contributed to rat and mouse histological analyses.

## Notes

### Competing Interest Statement

The authors have declared no competing interest.

## REFERENCES

1 CDC, N. Underlying Cause of Death 1999-2013 on CDC WONDER Online Database, released 2015. Data are from the Multiple Cause of Death Files, 1999-2013, as compiled from data provided by the 57 vital statistics jurisdictions through the Vital Statistics Cooperative Program.

2 Mozaffarian, D. et al. Heart disease and stroke statistics--2015 update: a report from the American Heart Association. Circulation 131, e29–322, doi:10.1161/CIR.0000000000000152 (2015).

3 Lian, X. et al. Robust cardiomyocyte differentiation from human pluripotent stem cells via temporal modulation of canonical Wnt signaling. Proc Natl Acad Sci U S A 109, E1848–1857, doi:10.1073/pnas.1200250109 (2012).

4 Burridge, P. W. et al. Chemically defined generation of human cardiomyocytes. Nat Methods 11, 855–860, doi:10.1038/nmeth.2999 (2014).

5 Prowse, A. B. et al. Transforming the promise of pluripotent stem cell-derived cardiomyocytes to a therapy: challenges and solutions for clinical trials. The Canadian journal of cardiology 30, 1335–1349, doi:10.1016/j.cjca.2014.08.005 (2014).

6 Vunjak-Novakovic, G. et al. Challenges in cardiac tissue engineering. Tissue engineering. Part B, Reviews 16, 169–187, doi:10.1089/ten.TEB.2009.0352 (2010).

7 Hirt, M. N., Hansen, A. & Eschenhagen, T. Cardiac tissue engineering: state of the art. Circulation research 114, 354–367, doi:10.1161/CIRCRESAHA.114.300522 (2014).

8 Barad, L., Schick, R., Zeevi-Levin, N., Itskovitz-Eldor, J. & Binah, O. Human embryonic stem cells vs human induced pluripotent stem cells for cardiac repair. The Canadian journal of cardiology 30, 1279–1287, doi:10.1016/j.cjca.2014.06.023 (2014).

9 Gao, L. et al. Large Cardiac Muscle Patches Engineered From Human Induced-Pluripotent Stem Cell–Derived Cardiac Cells Improve Recovery From Myocardial Infarction in Swine. Circulation 137, 1712–1730, doi:10.1161/circulationaha.117.030785 (2018).

10 Laflamme, M. A. et al. Cardiomyocytes derived from human embryonic stem cells in pro-survival factors enhance function of infarcted rat hearts. Nat Biotechnol 25, 1015–1024, doi:10.1038/nbt1327 (2007).

11 Stevens, K. R. et al. Physiological function and transplantation of scaffold-free and vascularized human cardiac muscle tissue. Proc Natl Acad Sci U S A 106, 16568–16573, doi:10.1073/pnas.0908381106 (2009).

12 Shiba, Y. et al. Human ES-cell-derived cardiomyocytes electrically couple and suppress arrhythmias in injured hearts. Nature 489, 322–325, doi:10.1038/nature11317 (2012).

13 Chong, J. J. et al. Human embryonic-stem-cell-derived cardiomyocytes regenerate non-human primate hearts. Nature 510, 273–277, doi:10.1038/nature13233 (2014).

14 Liu, Y. W. et al. Human embryonic stem cell-derived cardiomyocytes restore function in infarcted hearts of non-human primates. Nat Biotechnol 36, 597–605, doi:10.1038/nbt.4162 (2018).

15 Bargehr, J. et al. Epicardial cells derived from human embryonic stem cells augment cardiomyocyte-driven heart regeneration. Nat Biotechnol 37, 895–906, doi:10.1038/s41587-019-0197-9 (2019).

16 Silver, S. E., Barrs, R. W. & Mei, Y. Transplantation of Human Pluripotent Stem Cell-Derived Cardiomyocytes for Cardiac Regenerative Therapy. Front Cardiovasc Med 8, 707890, doi:10.3389/fcvm.2021.707890 (2021).

17 Weinberger, F. et al. Cardiac repair in guinea pigs with human engineered heart tissue from induced pluripotent stem cells. Sci Transl Med 8, 363ra148, doi:10.1126/scitranslmed.aaf8781 (2016).

18 Fan, C. et al. CHIR99021 and fibroblast growth factor 1 enhance the regenerative potency of human cardiac muscle patch after myocardial infarction in mice. J Mol Cell Cardiol 141, 1–10, doi:10.1016/j.yjmcc.2020.03.003 (2020).

19 Yeung, E. et al. Cardiac regeneration using human-induced pluripotent stem cell-derived biomaterial-free 3D-bioprinted cardiac patch in vivo. Journal of tissue engineering and regenerative medicine 13, 2031–2039, doi:10.1002/term.2954 (2019).

20 Kawaguchi, S. et al. Intramyocardial Transplantation of Human iPS Cell–Derived Cardiac Spheroids Improves Cardiac Function in Heart Failure Animals. JACC: Basic to Translational Science, doi:10.1016/j.jacbts.2020.11.017 (2021).

21 Schaefer, J. A., Guzman, P. A., Riemenschneider, S. B., Kamp, T. J. & Tranquillo, R. T. A cardiac patch from aligned microvessel and cardiomyocyte patches. Journal of tissue engineering and regenerative medicine 12, 546–556, doi:10.1002/term.2568 (2018).

22 Kobayashi, H. et al. Regeneration of Nonhuman Primate Hearts With Human Induced Pluripotent Stem Cell-Derived Cardiac Spheroids. Circulation 150, 611–621, doi:10.1161/CIRCULATIONAHA.123.064876 (2024).

23 Silver, S. E., Barrs, R. W. & Mei, Y. Transplantation of Human Pluripotent Stem Cell-Derived Cardiomyocytes for Cardiac Regenerative Therapy. Frontiers in Cardiovascular Medicine 8, doi:10.3389/fcvm.2021.707890 (2021).

24 Jebran, A. F. et al. Engineered heart muscle allografts for heart repair in primates and humans. Nature, doi:10.1038/s41586-024-08463-0 (2025).

25 Robey, T. E., Saiget, M. K., Reinecke, H. & Murry, C. E. Systems approaches to preventing transplanted cell death in cardiac repair. Journal of molecular and cellular cardiology 45, 567–581, doi:10.1016/j.yjmcc.2008.03.009 (2008).

26 Ogasawara, T. et al. Impact of extracellular matrix on engraftment and maturation of pluripotent stem cell-derived cardiomyocytes in a rat myocardial infarct model. Sci Rep 7, 8630, doi:10.1038/s41598-017-09217-x (2017).

27 Shiba, Y. et al. Allogeneic transplantation of iPS cell-derived cardiomyocytes regenerates primate hearts. Nature 538, 388–391, doi:10.1038/nature19815 (2016).

28 Ogle, B. M. et al. Distilling complexity to advance cardiac tissue engineering. Sci Transl Med 8, 342ps313, doi:10.1126/scitranslmed.aad2304 (2016).

29 Nunes, S. S. et al. Biowire: a platform for maturation of human pluripotent stem cell-derived cardiomyocytes. Nature methods 10, 781–787, doi:10.1038/nmeth.2524 (2013).

30 Mannhardt, I. et al. Human Engineered Heart Tissue: Analysis of Contractile Force. Stem cell reports 7, 29–42, doi:10.1016/j.stemcr.2016.04.011 (2016).

31 Gerbin, K. A., Yang, X., Murry, C. E. & Coulombe, K. L. Enhanced Electrical Integration of Engineered Human Myocardium via Intramyocardial versus Epicardial Delivery in Infarcted Rat Hearts. PloS one 10, e0131446, doi:10.1371/journal.pone.0131446 (2015).

32 Sekine, H. et al. Endothelial cell coculture within tissue-engineered cardiomyocyte sheets enhances neovascularization and improves cardiac function of ischemic hearts. Circulation 118, S145–152, doi:10.1161/CIRCULATIONAHA.107.757286 (2008).

33 Redd, M. A. et al. Patterned human microvascular grafts enable rapid vascularization and increase perfusion in infarcted rat hearts. Nature communications 10, 584, doi:10.1038/s41467-019-08388-7 (2019).

34 Richards, D. J. et al. Inspiration from heart development: Biomimetic development of functional human cardiac organoids. Biomaterials 142, 112–123, doi:10.1016/j.biomaterials.2017.07.021 (2017).

35 Hyams, N. A., Kerr, C. M., Arhontoulis, D. C., Ruddy, J. M. & Mei, Y. Improving human cardiac organoid design using transcriptomics. Scientific reports 14, doi:10.1038/s41598-024-61554-w (2024).

36 Tan, Y. et al. Nanowired human cardiac organoid transplantation enables highly efficient and effective recovery of infarcted hearts. Sci Adv 9, eadf2898, doi:10.1126/sciadv.adf2898 (2023).

37 Eschenhagen, T. & Weinberger, F. Challenges and perspectives of heart repair with pluripotent stem cell-derived cardiomyocytes. Nat Cardiovasc Res 3, 515–524, doi:10.1038/s44161-024-00472-6 (2024).

38 Koenig, A. et al. Missing self triggers NK cell-mediated chronic vascular rejection of solid organ transplants. Nat Commun 10, 5350, doi:10.1038/s41467-019-13113-5 (2019).

39 Gornalusse, G. G. et al. HLA-E-expressing pluripotent stem cells escape allogeneic responses and lysis by NK cells. Nat Biotechnol 35, 765–772, doi:10.1038/nbt.3860 (2017).

40 Deuse, T. et al. Hypoimmunogenic derivatives of induced pluripotent stem cells evade immune rejection in fully immunocompetent allogeneic recipients. Nat Biotechnol 37, 252–258, doi:10.1038/s41587-019-0016-3 (2019).

41 Hotta, A., Schrepfer, S. & Nagy, A. Genetically engineered hypoimmunogenic cell therapy. Nature Reviews Bioengineering, doi:10.1038/s44222-024-00219-9 (2024).

42 Han, X. et al. Generation of hypoimmunogenic human pluripotent stem cells. Proc Natl Acad Sci U S A 116, 10441–10446, doi:10.1073/pnas.1902566116 (2019).

43 Hu, X. et al. Hypoimmune induced pluripotent stem cells survive long term in fully immunocompetent, allogeneic rhesus macaques. Nat Biotechnol, doi:10.1038/s41587-023-01784-x (2023).

44 Hu, X. et al. Human hypoimmune primary pancreatic islets avoid rejection and autoimmunity and alleviate diabetes in allogeneic humanized mice. Sci Transl Med 15, eadg5794, doi:10.1126/scitranslmed.adg5794 (2023).

45 Hu, X. et al. Hypoimmune islets achieve insulin independence after allogeneic transplantation in a fully immunocompetent non-human primate. Cell stem cell 31, 334–340 e335, doi:10.1016/j.stem.2024.02.001 (2024).

46 Strong, R. K. et al. HLA-E Allelic Variants. Journal of Biological Chemistry 278, 5082–5090, doi:10.1074/jbc.m208268200 (2003).

47 Rouas-Freiss, N., Marchal, R. E., Kirszenbaum, M., Dausset, J. & Carosella, E. D. The alpha1 domain of HLA-G1 and HLA-G2 inhibits cytotoxicity induced by natural killer cells: is HLA-G the public ligand for natural killer cell inhibitory receptors? Proceedings of the National Academy of Sciences of the United States of America 94, 5249–5254, doi:10.1073/pnas.94.10.5249 (1997).

48 Naji, A. et al. Binding of HLA-G to ITIM-Bearing Ig-like Transcript 2 Receptor Suppresses B Cell Responses. The Journal of Immunology 192, 1536–1546, doi:10.4049/jimmunol.1300438 (2014).

49 Pazmany, L. et al. Protection from Natural Killer Cell-Mediated Lysis by HLA-G Expression on Target Cells. Science 274, 792–795, doi:10.1126/science.274.5288.792 (1996).

50 Dumont, C. et al. CD8+PD-1–ILT2+ T Cells Are an Intratumoral Cytotoxic Population Selectively Inhibited by the Immune-Checkpoint HLA-G. Cancer Immunology Research 7, 1619–1632, doi:10.1158/2326-6066.cir-18-0764 (2019).

51 Llano, M. et al. HLA-E-bound peptides influence recognition by inhibitory and triggering CD94/NKG2 receptors: preferential response to an HLA-G-derived nonamer. European Journal of Immunology 28, 2854–2863, doi:10.1002/(sici)1521-4141(199809)28:09<2854::aid-immu2854>3.0.co;2-w (1998).

52 André, P. et al. Anti-NKG2A mAb Is a Checkpoint Inhibitor that Promotes Anti-tumor Immunity by Unleashing Both T and NK Cells. Cell 175, 1731–1743.e1713, doi:10.1016/j.cell.2018.10.014 (2018).

53 Carosella, E. D., Rouas-Freiss, N., Roux, D. T.-L., Moreau, P. & LeMaoult, J. in Advances in Immunology 33–144 (Elsevier, 2015).

54 Clements, C. S. et al. Crystal structure of HLA-G: a nonclassical MHC class I molecule expressed at the fetal-maternal interface. Proceedings of the National Academy of Sciences of the United States of America 102, 3360–3365, doi:10.1073/pnas.0409676102 (2005).

55 Ferreira, L. M. R., Meissner, T. B., Tilburgs, T. & Strominger, J. L. HLA-G: At the Interface of Maternal-Fetal Tolerance. Trends Immunol 38, 272–286, doi:10.1016/j.it.2017.01.009 (2017).

56 Ishitani, A. et al. Protein Expression and Peptide Binding Suggest Unique and Interacting Functional Roles for HLA-E, F, and G in Maternal-Placental Immune Recognition. The Journal of Immunology 171, 1376–1384, doi:10.4049/jimmunol.171.3.1376 (2003).

57 Haapaniemi, E., Botla, S., Persson, J., Schmierer, B. & Taipale, J. CRISPR–Cas9 genome editing induces a p53-mediated DNA damage response. Nature Medicine 24, 927–930, doi:10.1038/s41591-018-0049-z (2018).

58 Ihry, R. J. et al. p53 inhibits CRISPR–Cas9 engineering in human pluripotent stem cells. Nature Medicine 24, 939–946, doi:10.1038/s41591-018-0050-6 (2018).

59 Mali, P. et al. RNA-guided human genome engineering via Cas9. Science (New York, N.Y.) 339, 823–826, doi:10.1126/science.1232033 (2013).

60 Horowitz, A. et al. Genetic and environmental determinants of human NK cell diversity revealed by mass cytometry. Sci Transl Med 5, 208ra145, doi:10.1126/scitranslmed.3006702 (2013).

61 Kawaguchi, S. et al. Intamyocardial transplantation of human iPS cell-derived cardiac spheroids improves cardiac function in heart failure animals. J Am Coll Cardiol Basic Trans Science (2021).

62 Giacomelli, E. et al. Human-iPSC-Derived Cardiac Stromal Cells Enhance Maturation in 3D Cardiac Microtissues and Reveal Non-cardiomyocyte Contributions to Heart Disease. Cell Stem Cell 26, 862–879 e811, doi:10.1016/j.stem.2020.05.004 (2020).

63 Campostrini, G. et al. Generation, functional analysis and applications of isogenic three-dimensional self-aggregating cardiac microtissues from human pluripotent stem cells. Nat Protoc 16, 2213–2256, doi:10.1038/s41596-021-00497-2 (2021).

64 Karbassi, E. et al. Cardiomyocyte maturation: advances in knowledge and implications for regenerative medicine. Nature reviews. Cardiology 17, 341–359, doi:10.1038/s41569-019-0331-x (2020).

65 Burchfield, J. S., Xie, M. & Hill, J. A. Pathological ventricular remodeling: mechanisms: part 1 of 2. Circulation 128, 388–400, doi:10.1161/CIRCULATIONAHA.113.001878 (2013).

66 Hu, X. et al. Hypoimmune induced pluripotent stem cells survive long term in fully immunocompetent, allogeneic rhesus macaques. Nat Biotechnol 42, 413–423, doi:10.1038/s41587-023-01784-x (2024).

67 Dubois, N. C. et al. SIRPA is a specific cell-surface marker for isolating cardiomyocytes derived from human pluripotent stem cells. Nat Biotechnol 29, 1011–1018, doi:10.1038/nbt.2005 (2011).

68 Haideri, T. et al. Robust genome editing via modRNA-based Cas9 or base editor in human pluripotent stem cells. Cell Rep Methods 2, 100290, doi:10.1016/j.crmeth.2022.100290 (2022).

69 He, W. et al. Intracellular trafficking of HLA-E and its regulation. J Exp Med 220, doi:10.1084/jem.20221941 (2023).

70 Blazeski, A., Garcia-Cardena, G. & Kamm, R. D. Advancing Cardiac Organoid Engineering Through Application of Biophysical Forces. IEEE Rev Biomed Eng **PP**, doi:10.1109/RBME.2024.3514378 (2024).

71 Drakhlis, L. et al. Human heart-forming organoids recapitulate early heart and foregut development. Nature biotechnology, doi:10.1038/s41587-021-00815-9 (2021).

72 Hofbauer, P. et al. Cardioids reveal self-organizing principles of human cardiogenesis. Cell, doi:10.1016/j.cell.2021.04.034 (2021).

73 Richards, D. J. et al. Human cardiac organoids for the modelling of myocardial infarction and drug cardiotoxicity. Nat Biomed Eng 4, 446–462, doi:10.1038/s41551-020-0539-4 (2020).

74 Arhontoulis, D. C. et al. Human cardiac organoids to model COVID-19 cytokine storm induced cardiac injuries. J Tissue Eng Regen Med 16, 799–811, doi:10.1002/term.3327 (2022).

75 Lian, X. et al. A small molecule inhibitor of SRC family kinases promotes simple epithelial differentiation of human pluripotent stem cells. PloS one 8, e60016–e60016, doi:10.1371/journal.pone.0060016 (2013).

76 Bao, X. et al. Chemically-defined albumin-free differentiation of human pluripotent stem cells to endothelial progenitor cells. Stem Cell Res 15, 122–129, doi:10.1016/j.scr.2015.05.004 (2015).

77 Young, L., Sung, J., Stacey, G. & Masters, J. R. Detection of Mycoplasma in cell cultures. Nature Protocols 5, 929–934, doi:10.1038/nprot.2010.43 (2010).

78 Lian, X. et al. Chemically defined, albumin-free human cardiomyocyte generation. Nat Methods 12, 595–596, doi:10.1038/nmeth.3448 (2015).

79 Lian, X. et al. Directed cardiomyocyte differentiation from human pluripotent stem cells by modulating Wnt/beta-catenin signaling under fully defined conditions. Nat Protoc 8, 162–175, doi:10.1038/nprot.2012.150 (2013).

80 Hansen, T., Yu, Y. Y. L. & Fremont, D. H. Preparation of Stable Single-Chain Trimers Engineered with Peptide, β2 Microglobulin, and MHC Heavy Chain. Current Protocols in Immunology 87, doi:10.1002/0471142735.im1705s87 (2009).

81 Kao, T. et al. GAPTrap: A Simple Expression System for Pluripotent Stem Cells and Their Derivatives. Stem cell reports 7, 518–526, doi:10.1016/j.stemcr.2016.07.015 (2016).

82 Karimi, M. A. et al. Measuring cytotoxicity by bioluminescence imaging outperforms the standard chromium-51 release assay. PloS one 9, e89357–e89357, doi:10.1371/journal.pone.0089357 (2014).

83 Shultz, L. D. et al. Subcapsular transplantation of tissue in the kidney. Cold Spring Harb Protoc 2014, 737–740, doi:10.1101/pdb.prot078089 (2014).

84 Szot, G. L., Koudria, P. & Bluestone, J. A. Transplantation of Pancreatic Islets Into the Kidney Capsule of Diabetic Mice. Journal of Visualized Experiments, doi:10.3791/404-v (2007).

85 Kerr, C. M. et al. Decellularized heart extracellular matrix alleviates activation of hiPSC-derived cardiac fibroblasts. Bioact Mater 31, 463–474, doi:10.1016/j.bioactmat.2023.08.023 (2024).

86 Tohyama, S. et al. Distinct metabolic flow enables large-scale purification of mouse and human pluripotent stem cell-derived cardiomyocytes. Cell Stem Cell 12, 127–137, doi:10.1016/j.stem.2012.09.013 (2013).

87 Maas, R. G. C. et al. Massive expansion and cryopreservation of functional human induced pluripotent stem cell-derived cardiomyocytes. STAR Protoc 2, 100334, doi:10.1016/j.xpro.2021.100334 (2021).

88 Buikema, J. W. et al. Wnt Activation and Reduced Cell-Cell Contact Synergistically Induce Massive Expansion of Functional Human iPSC-Derived Cardiomyocytes. Cell Stem Cell 27, 50–63 e55, doi:10.1016/j.stem.2020.06.001 (2020).

89 Bao, X. et al. Long-term self-renewing human epicardial cells generated from pluripotent stem cells under defined xeno-free conditions. Nat Biomed Eng 1, doi:10.1038/s41551-016-0003 (2016).

90 Bao, X. et al. Directed differentiation and long-term maintenance of epicardial cells derived from human pluripotent stem cells under fully defined conditions. Nat Protoc 12, 1890–1900, doi:10.1038/nprot.2017.080 (2017).

91 Jin, Y. et al. Three-dimensional heart extracellular matrix enhances chemically induced direct cardiac reprogramming. Sci Adv 8, eabn5768, doi:10.1126/sciadv.abn5768 (2022).

92 Cho, H. et al. iPSC-derived endothelial cell response to hypoxia via SDF1a/CXCR4 axis facilitates incorporation to revascularize ischemic retina. JCI Insight 5, doi:10.1172/jci.insight.131828 (2020).

93 Smith, Q. et al. Differential HDAC6 Activity Modulates Ciliogenesis and Subsequent Mechanosensing of Endothelial Cells Derived from Pluripotent Stem Cells. Cell Rep 24, 895–908 e896, doi:10.1016/j.celrep.2018.06.083 (2018).

94 Bao, X., Lian, X. & Palecek, S. P. Directed Endothelial Progenitor Differentiation from Human Pluripotent Stem Cells Via Wnt Activation Under Defined Conditions. Methods Mol Biol 1481, 183–196, doi:10.1007/978-1-4939-6393-5_17 (2016).

95 Nishihara, H. et al. Advancing human induced pluripotent stem cell-derived blood-brain barrier models for studying immune cell interactions. FASEB J 34, 16693–16715, doi:10.1096/fj.202001507RR (2020).

96 Faal, T. et al. Induction of Mesoderm and Neural Crest-Derived Pericytes from Human Pluripotent Stem Cells to Study Blood-Brain Barrier Interactions. Stem Cell Reports 12, 451–460, doi:10.1016/j.stemcr.2019.01.005 (2019).

97 Zimmerman, C. M., Robino, R. A., Cochrane, R. W., Dominguez, M. D. & Ferreira, L. M. R. Redirecting Human Conventional and Regulatory T Cells Using Chimeric Antigen Receptors. Methods Mol Biol 2748, 201–241, doi:10.1007/978-1-0716-3593-3_15 (2024).

98 Barra, J. M. et al. Combinatorial genetic engineering strategy for immune protection of stem cell-derived beta cells by chimeric antigen receptor regulatory T cells. Cell Rep 43, 114994, doi:10.1016/j.celrep.2024.114994 (2024).

