## Supplementary Fig. for "Hypoimmunogenic hPSC-derived cardiac organoids for immune evasion and heart repair"

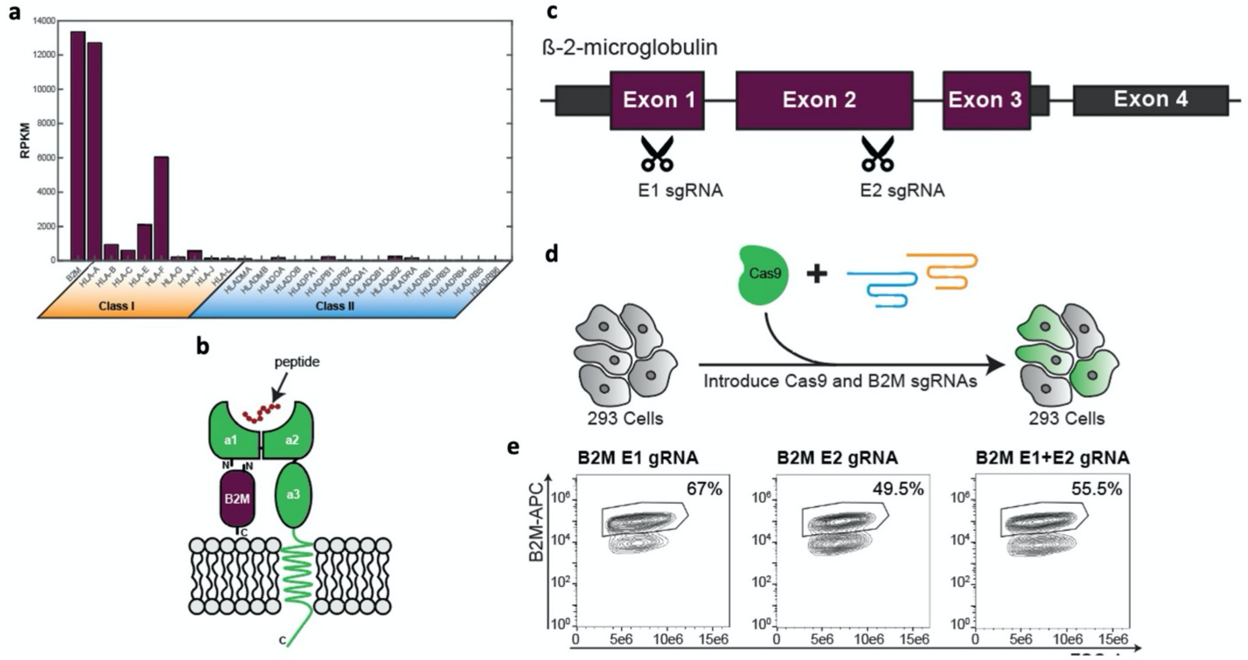


**Supplementary Fig. 1: Design strategy for generation of B2M KO hPSCs. a**, Relative expression for MHC molecules from bulk RNA sequencing analysis of day 60 hPSC derived CMs. **b,** Schematic illustrating the structure of class I HLA molecules, illustrating the specific HLA heavy chain noncovalently complexed to both a conserved B2M and a presented antigen peptide. **c,** Schematic illustrating the 4 exons in B2M and the locations in the B2M gene where sgRNAs targeted Exon 1 (E1 sgRNA) or Exon 2 (E2 sgRNA). **d,** Experimental schematic for the generation of a B2M KO with HEK 293 cells, illustrating the lipofection of cells with a plasmid encoding for Cas9, GFP, and sgRNAs. **e,** Flow cytometry analysis of B2M expression for HEK 293 cells one week after transfection.

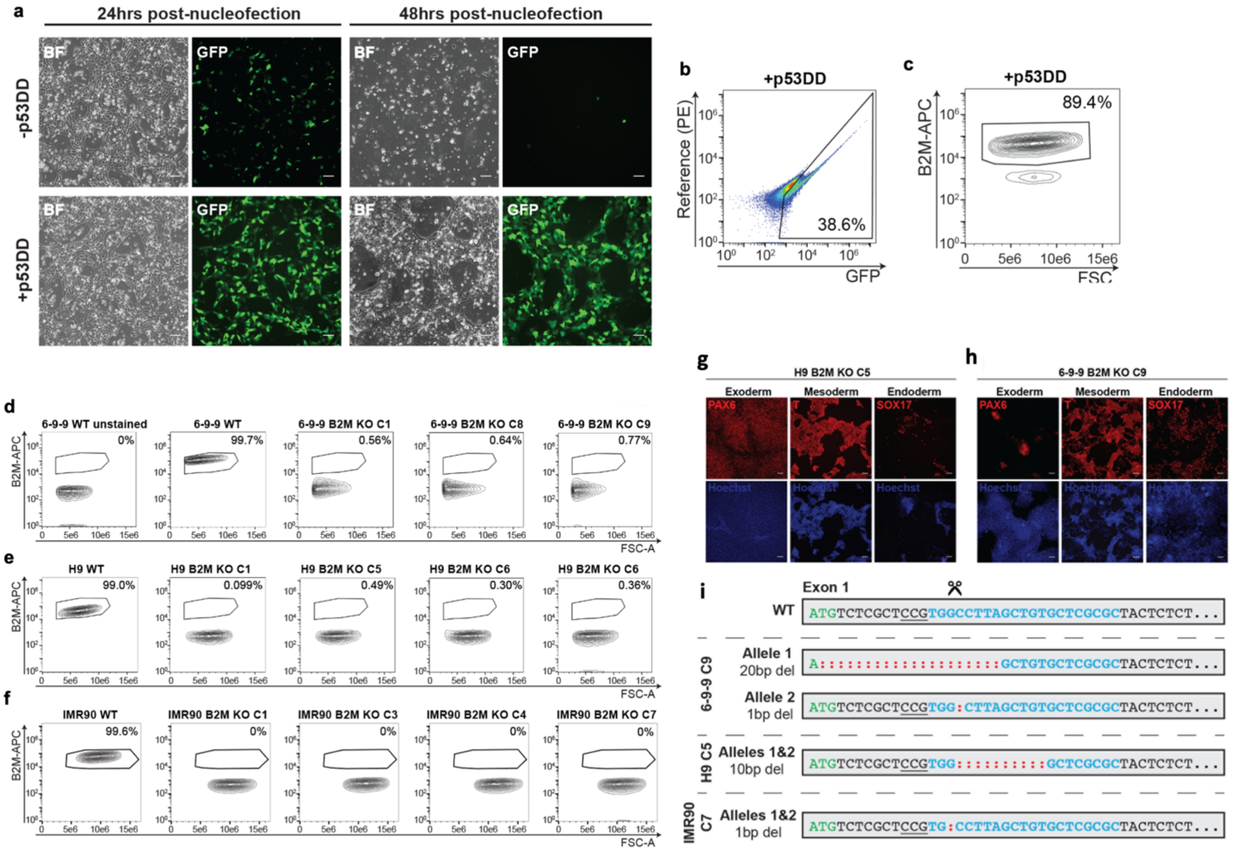


**Supplementary Fig. 2: Generation and characterization of B2M KO hPSCs. a**, Brightfield and GFP images of B2M KO cells transfected with p53DD (bottom) and without (top) 24 and 48 hours after nucleofection. Scale bars are 100 µm. **b,** Flow cytometry quantification of the GFP positive percentage of p53DD treated cells 48 hours after nucleofection. **c,** Flow cytometry of analysis of B2M expression for a mixed population of cells treated with p53DD one week after nucleofection. **d-f**, Flow cytometry-based identification of successful B2M KO clones derived from (**d**) 6-9-9, (**e**) H9, and (**f**) IMR90 cells. Wild type cells from each cell line and unstained control are also shown. **g-h**, Immunostaining validating differentiation of (**g**) H9 and (**h**) 6-9-9 B2M KO into to all three germ layers. Scale bars are 100 µm. **i**, Sanger sequencing of WT and KO alleles revealing frame shift mutations and premature stop codons. Green indicates start codon. Bolded blue indicates sgRNA sequence. Underline indicates the protospacer adjacent motif. Red colons indicate deleted base pairs.

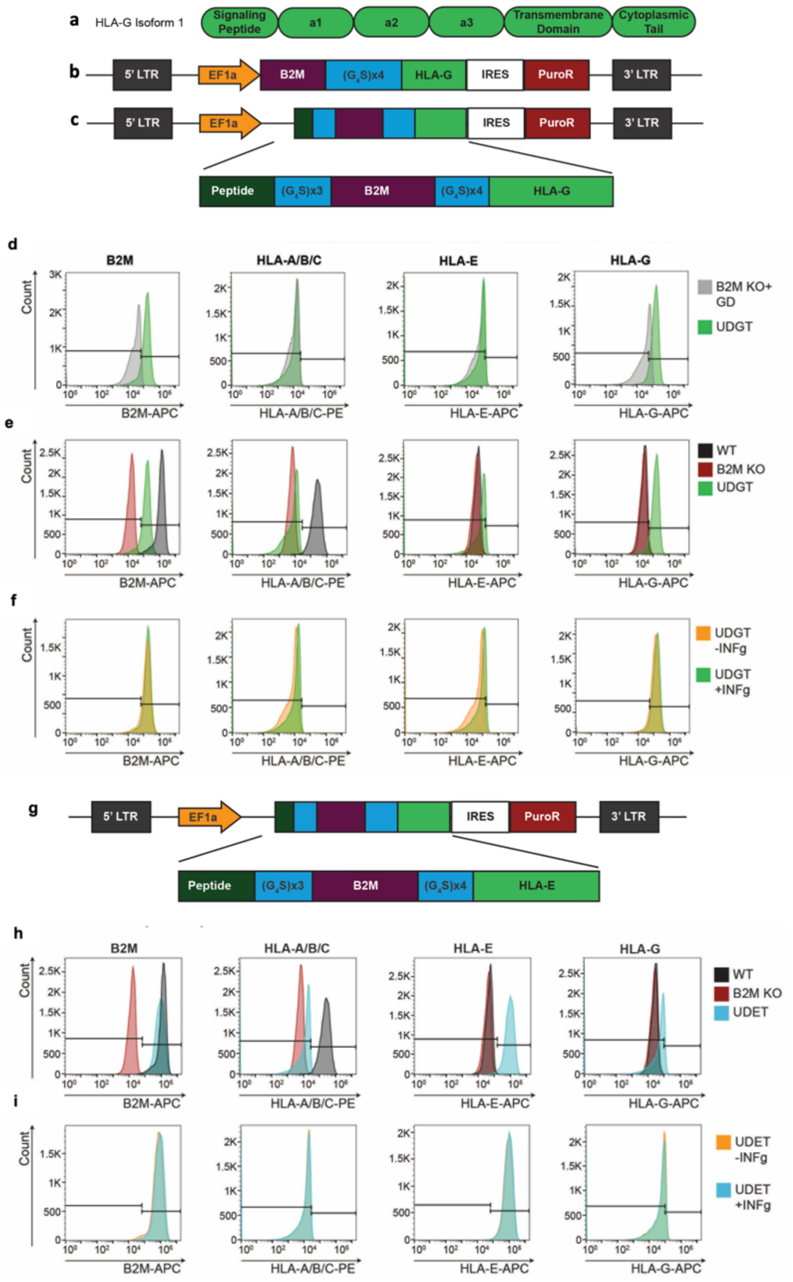


**Supplementary Fig. 3: Lentiviral delivery of HLA-G and HLA-E trimers promote robust engineered surface expression within undifferentiated hPSCs. a**, Schematic showing HLA-G isoform 1 components. **b-c,** Plasmid maps for (**b**) HLA-G dimer construct sequence containing B2M and HLA-G connected by a flexible non-cleavable linker and (**c**) HLA-G trimer construct sequence, illustrating constitutive promotors with a puromycin resistance gene. **d,** Comparative live cell flow cytometry analysis of HLA expression for B2M KO+GD and UDGT hPSCs. **e**, Flow cytometry analysis of HLA expression in WT (black), B2M KO (red), and UDGT (green) hPSCs. **f**, Comparative live cell flow cytometry analysis of HLA expression for UDGT hPSCs treated with or without INF$\gamma$. **g**, Plasmid map for HLA-E trimer construct sequence introduced via lentivirus. **h,** Flow cytometry analysis of HLA expression in WT (black), B2M KO (red), and HLA-E trimer (blue) hPSCs. **i**, Flow cytometry analysis of HLA expression in H9 B2M KO hPSCs transduced with HLA-E trimer treated with or without INF$\gamma$.

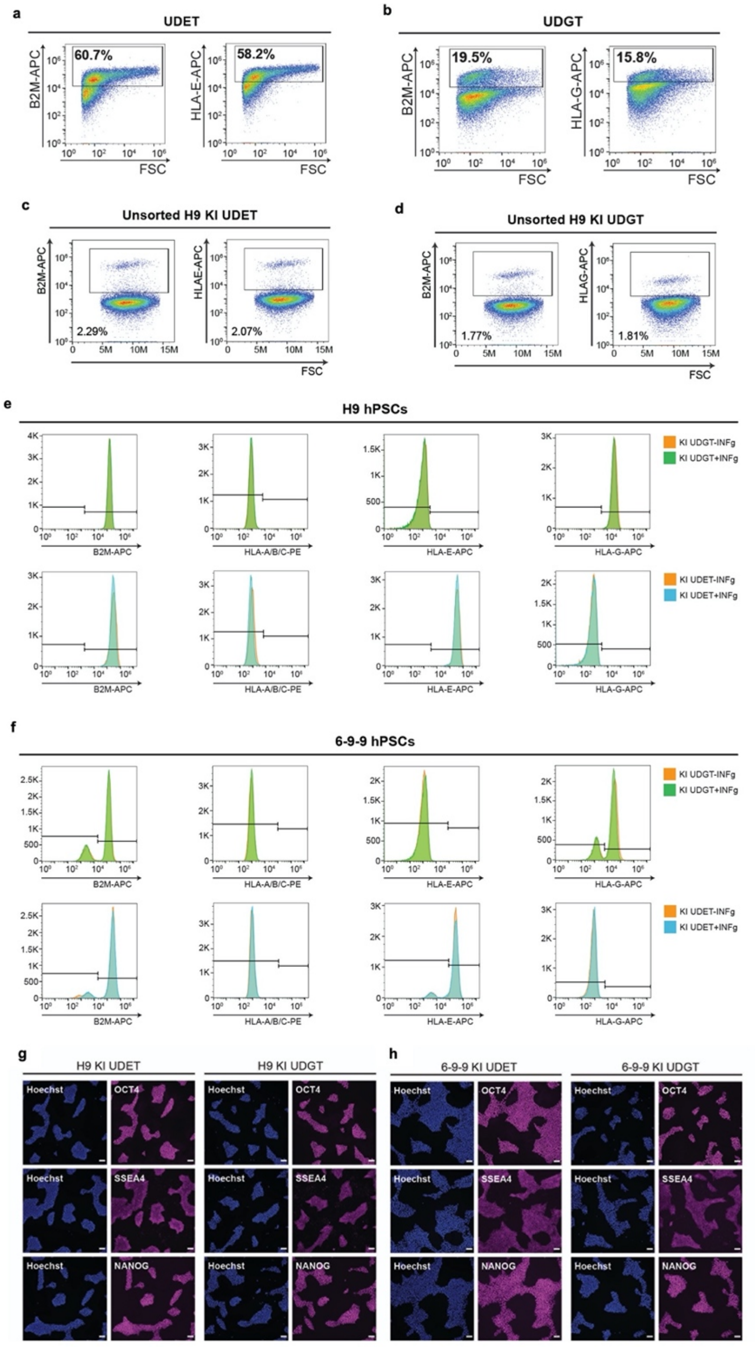


**Supplementary Fig. 4: AAVS1 safe harbor KI of HLA trimer construct is required for stable surface expression after differentiation. a-b**, Flow cytometry analysis of lentiviral engineered HLA expression in D8 cardiac progenitor cells derived from (**a**) H9 UDET and (**b**) H9 UDGT. **c-d**, Flow cytometry analysis of B2M and (**c**) HLA-E or (**d**) HLA-G to determine Cas9 mediated knock in efficiency seven days after nucleofection. **e-f**, Flow cytometry analysis of HLA expression for (**e**) 6-9-9 and (**f**) H9 KI UDGT (top) and KI UDET (bottom) hPSCs treated with or without INF$\gamma$. **g-h**, Immunostaining for pluripotency genes for (**g**) H9 KI hPSCs and (**h**) 6-9-9 KI hPSCs.

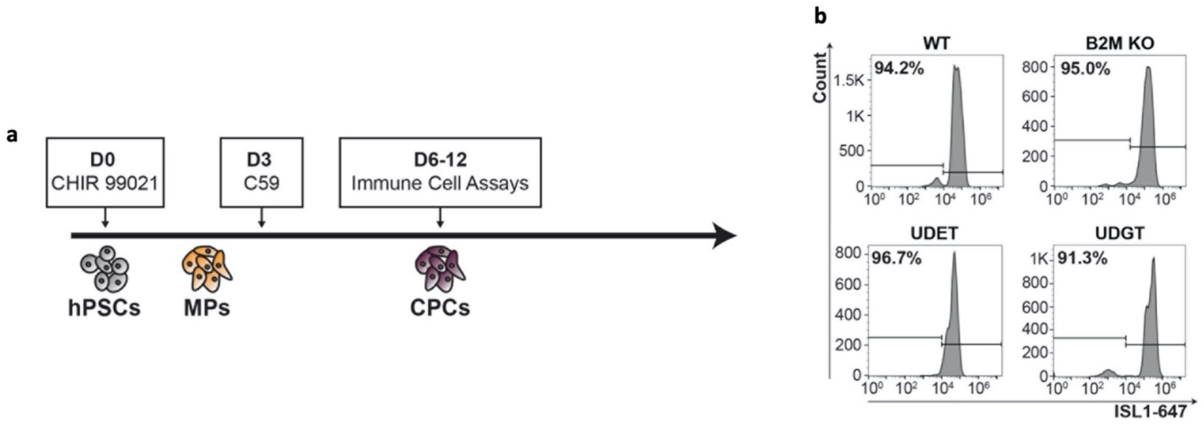


**Supplementary Fig. 5: Development and characterization of hPSC-CPCs. a**, Schematic of CPC differentiation. **b**, Analysis of day 6 cells for expression of cardiac progenitor marker ISL1.

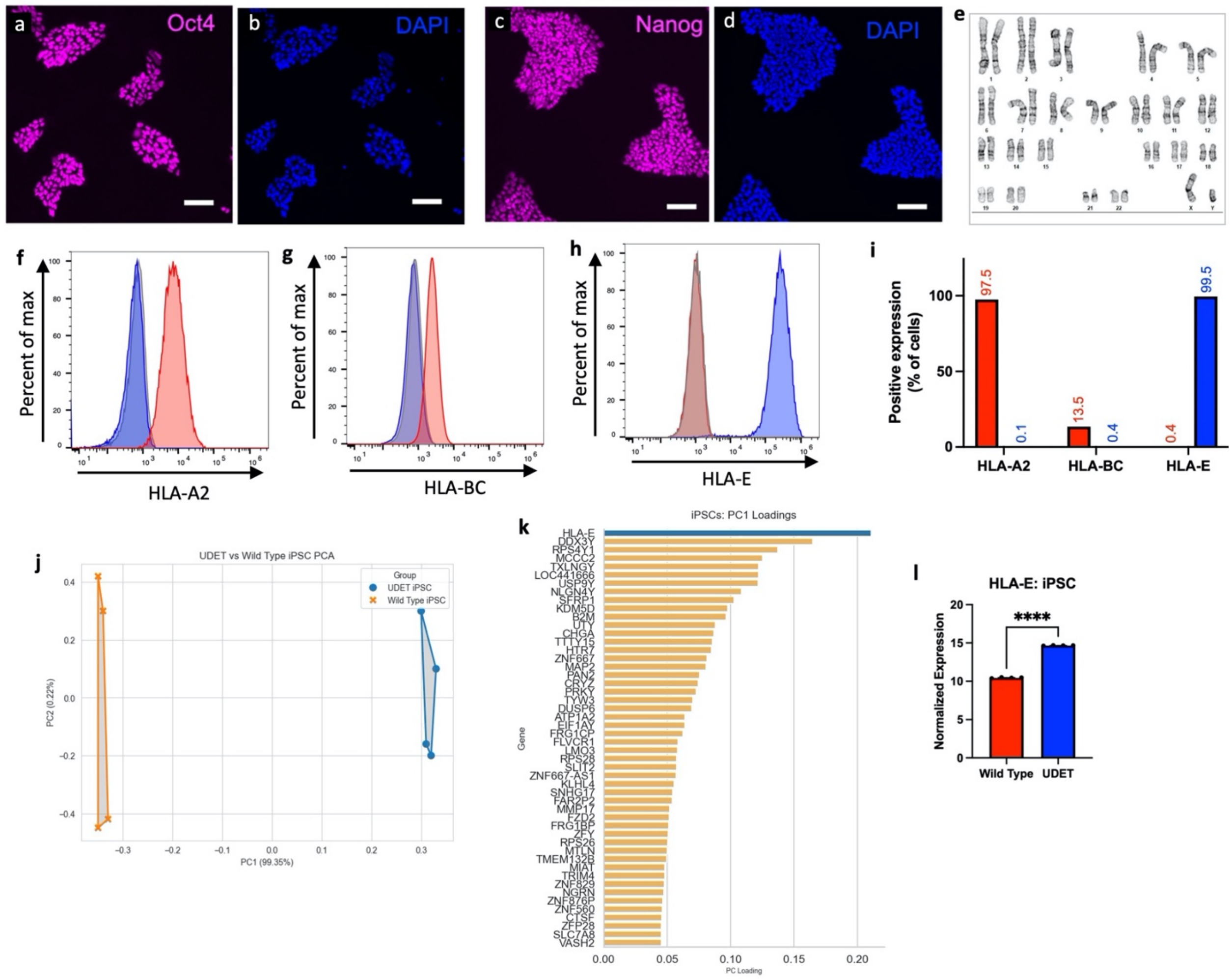


**Supplementary Fig. 6: Characterization of WT and KI UDET hPSCs generated from the 19-9-11 hiPSC line. a-d**, Immunostaining of pluripotency markers **(a,b)** Oct4 (purple) and (**c,d**) Nanog (purple), counterstained with DAPI (blue). **e,** Normal karyotype of 19-9-11 KI UDET hPSCs ~30 passages after initial gene edit. **f-h,** Flow cytometry assessing the expression of (**f**) HLA-A2, (**g**) HLA-BC, and (**h**) HLA-E in wild-type (red) and UDET (blue) hPSCs and their corresponding isotype controls (gray).**i**, Bar chart of flow cytometry data showing expression of HLA-A2, HLA-BC, and HLA-E as percent positive expression. **j,** PCA detailing the separation between WT hPSCs and KI UDET hPSCs. **k**, Positive PC1 loadings used in the generation of the PC plot in **j**. **l**, Differential expression of HLA-E in WT and KI UDET hPSCs.

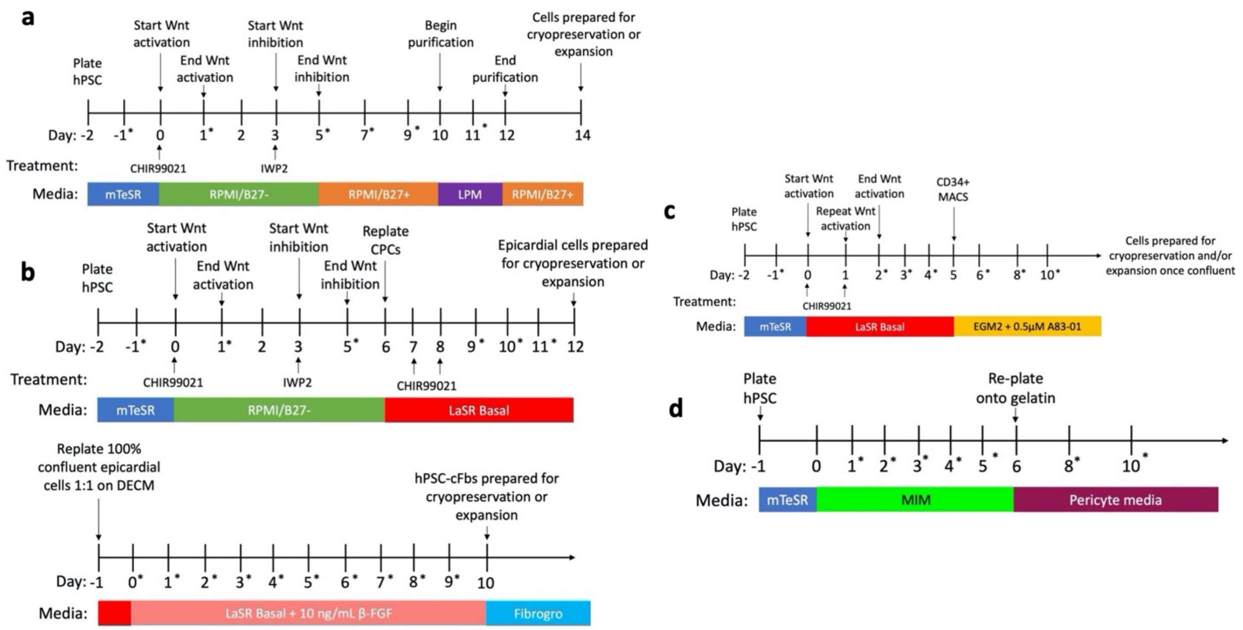


**Supplementary Fig. 7: Timelines for differentiation of hPSCs. a-d,** Steps as well as timing for the differentiation of (**a**) hPSC-CMs, (**b**) hPSC-cFbs derived from epicardial cells, (**c**) hPSC-ECs, and (**d**) hPSC-PCs.

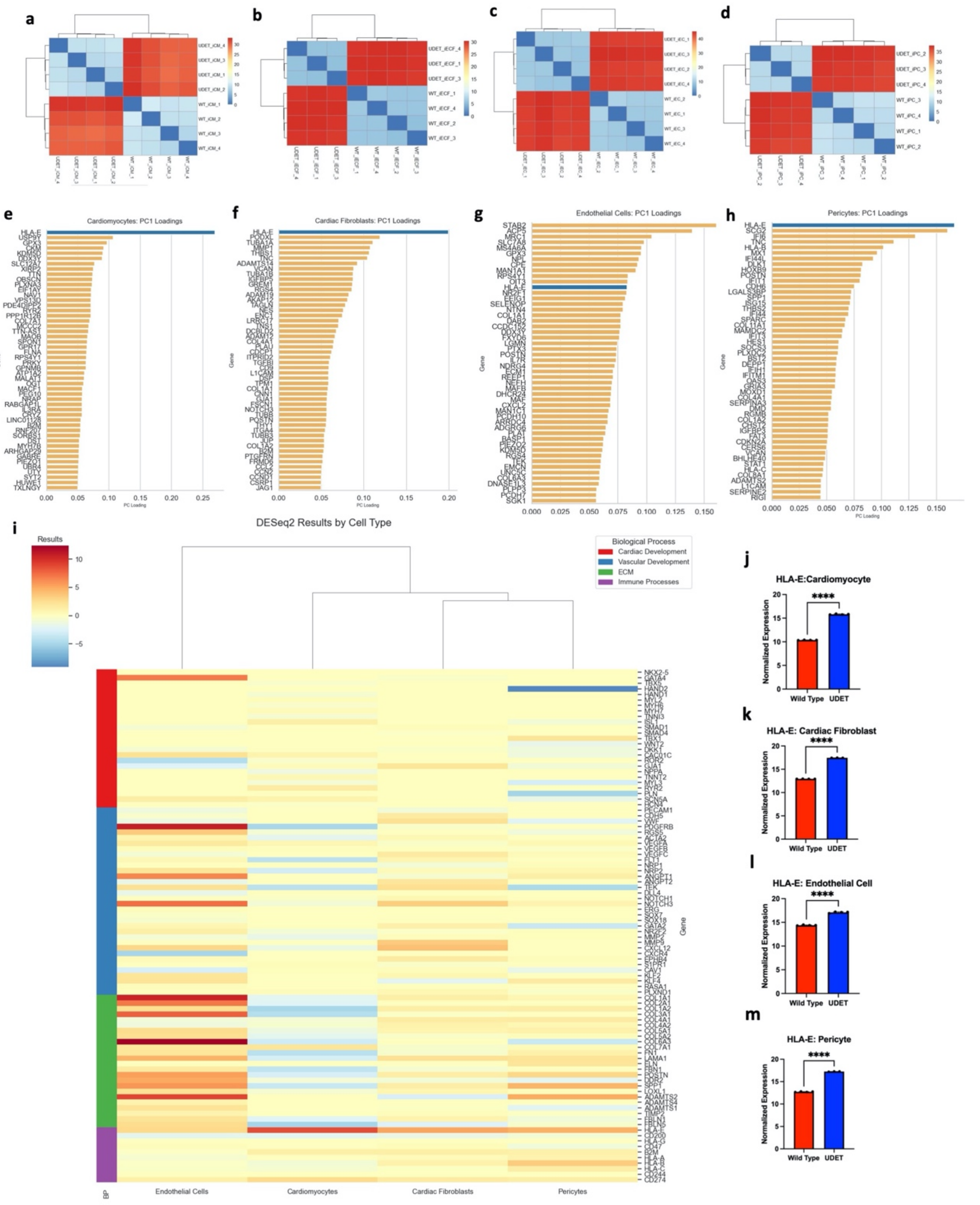


**Supplementary Fig. 8: Bulk RNA sequencing characterization of WT and KI UDET hPSC-derived cardiac cell types. a,** Sample distance heatmaps, representing general transcriptomic distance between (**a**) hPSC-CMs, (**b**) hPSC-cFbs, (**c**) hPSC-ECs, (**d**) hPSC-PCs. **e-h**, Positive PC1 loadings used in the generation of the PC plots in **Fig. 3n-q** for (**e**) hPSC-CMs in **Fig. 3n**, (**f**) hPSC-cFbs in **Fig. 3o**, (**g**) hPSC-ECs in **Fig. 3p**, and (**h**) hPSC-PCs in **Fig. 3q**. **i**, Heat map of differential expressed genes between WT and KI UDET hPSC-derived cells, highlighting genes involved in cardiac development, vascular development, ECM production, and immune processes for each hPSC-derived cardiac cell type. **j-m**, Differential expression of HLA-E in WT and UDET (**j**) hPSC-CMs, (**k**) hPSC-cFbs, (**l**) hPSC-ECs, (**m**) hPSC-PCs.

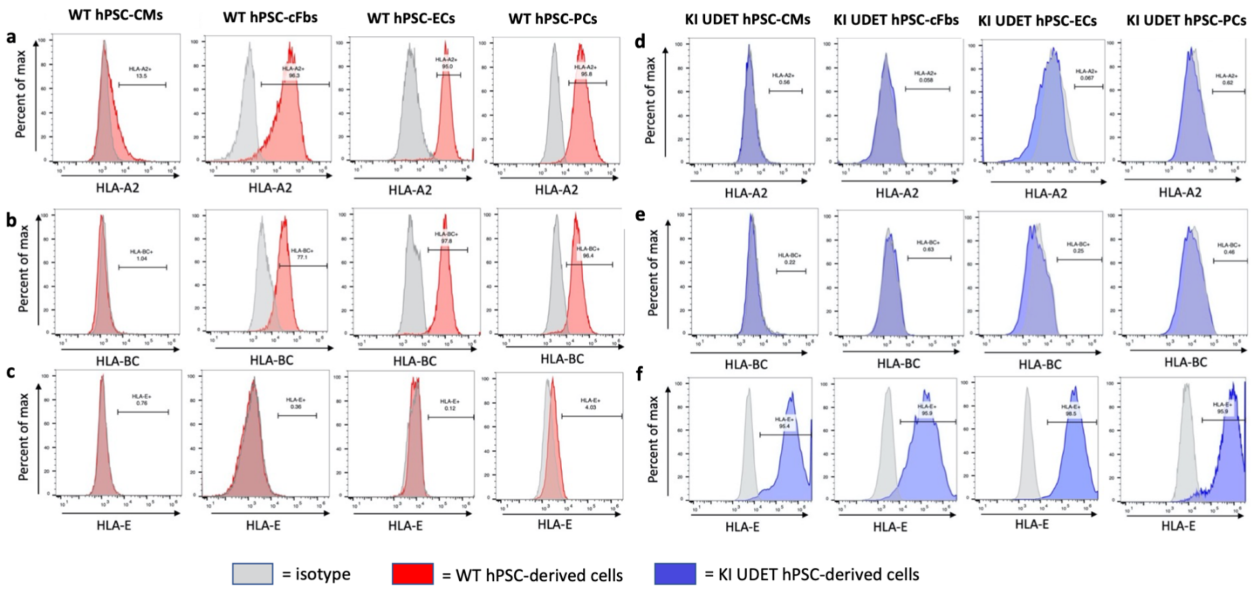


**Supplementary Fig. 9: Flow cytometry analysis of HLA expression in WT and KI UDET hPSC-derived cells**. **a-f,** Flow cytometry data used to create bar charts in **Fig. 4a-d** corresponding to the expression of (**a,d**) HLA-A2, (**b,e**) HLA-BC, and (**c,f**) HLA-E in (**a-c**) WT hPSC-derived and (**d-f**) UDET hPSC-derived hPSC-CMs, hPSC-cFbs, hPSC-ECs, and hPSC-PCs.

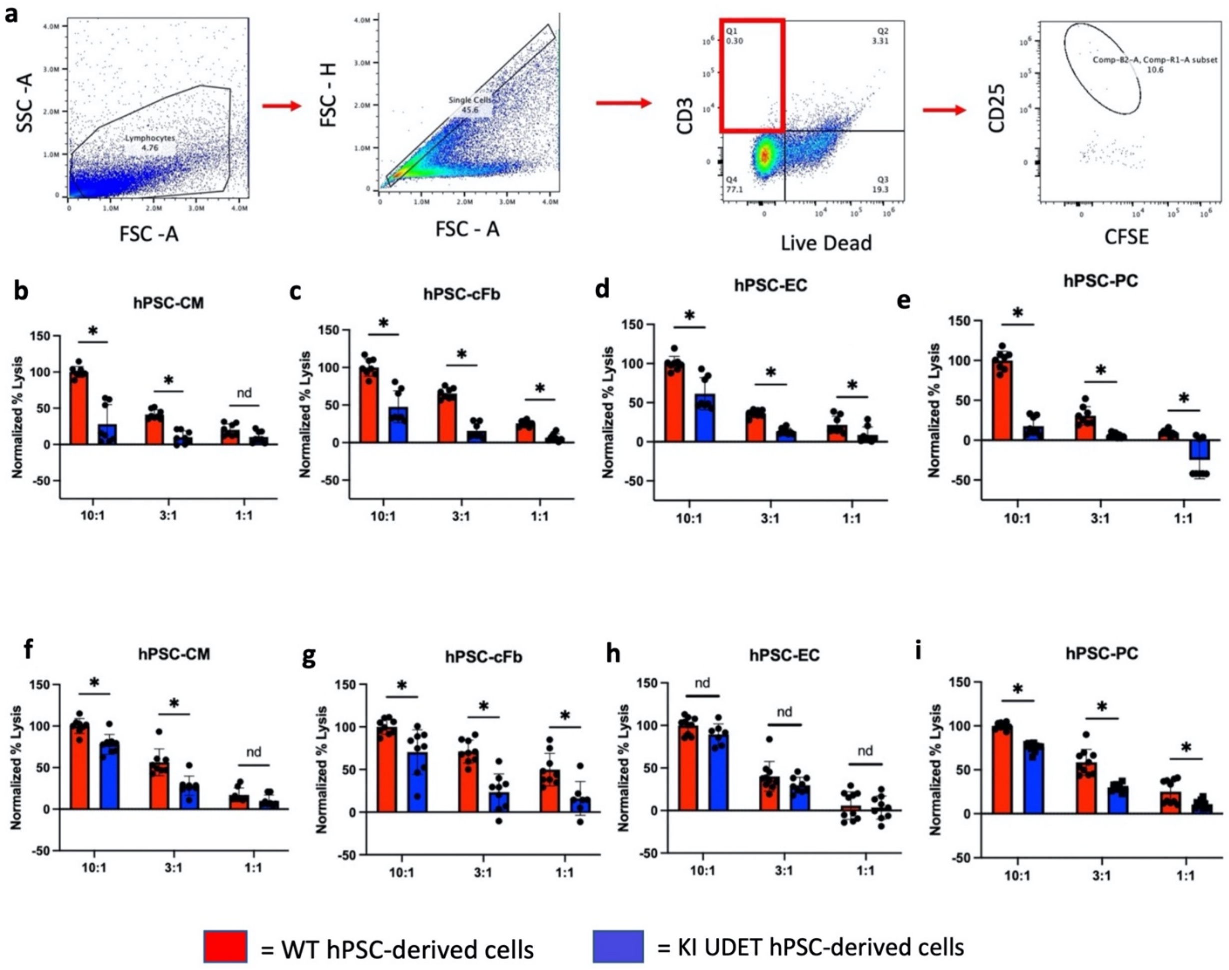


**Supplementary Fig. 10: Flow cytometry analysis of HLA-A2 negative CD8+ T cells cocultured with wild-type EB-derived cells and compiled CD8+ T cell and NK cell cytotoxicity assays normalized across 2 donors. a,** Gating strategy for CFSE-loaded T cells (left to right), showing T cell activation with negative expression of CSFE and positive expression of CD25. **b-e**, Normalized percent lysis for each donor quantified from LDH assay following exposure of activated CD8+ T cells to WT (red) and KI UDET (blue) (**b**) hPSC-CMs, (**c**) hPSC-cFbs, (**d**) hPSC-ECs, and (**e**) hPSC-PCs. **f-i**, Normalized percent lysis for each donor quantified from LDH assay following exposure of NK cells to WT (red) and KI UDET (blue) (**f**) hPSC-CMs, (**g**) hPSC-cFbs, (**h**) hPSC-ECs, and (**i**) hPSC-PCs.

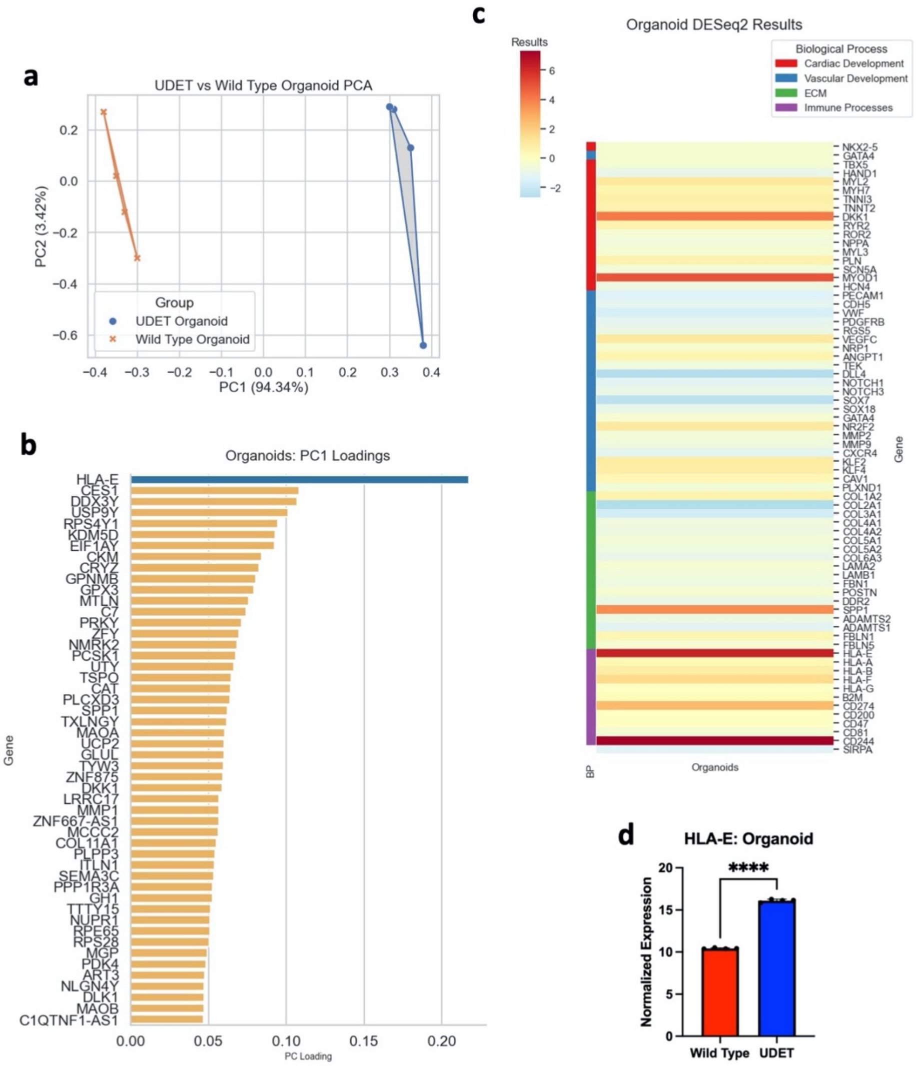


**Supplementary Fig. 11: Bulk RNA sequencing characterization of WT and KI UDET hPSC-derived cardiac organoids.** **a**, PCA detailing the separation between WT hPSCs and KI UDET hPSC-derived cardiac organoids. **b**, Positive PC1 loadings used in the generation of the PC plot in **a**. **c**, Heat map of differential expressed genes between WT and KI UDET hPSC-derived cardiac organoids, highlighting genes involved in cardiac development, vascular development, ECM production, and immune processes. **d**, Differential expression of HLA-E in WT and KI UDET hPSC-derived cardiac organoids.

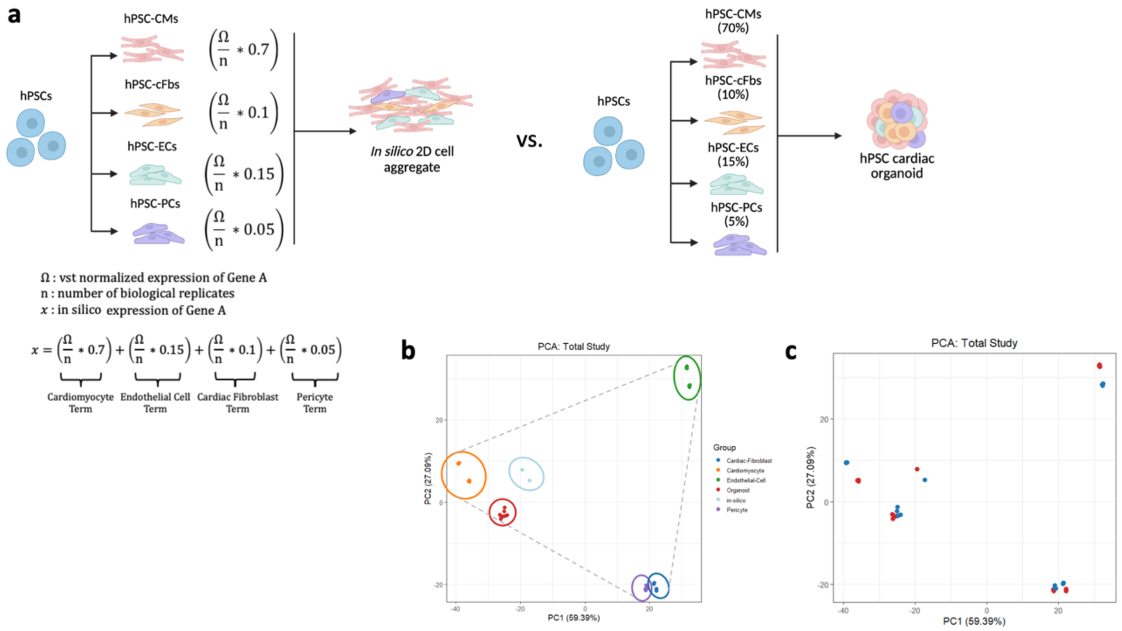


**Supplementary Fig. 12: Development of *in silico* cardiac organoids. a**, Development of *in silico* 2D aggregates (left), combining whole-transcriptome data from 2D cells based on the proportion of each cell type in hPSC-derived cardiac organoids (right) **b,** PCA detailing 2D hPSC-derived cell types, *in silico* 2D aggregates, and hPSC-derived cardiac organoids, outlining organoid and *in silico* 2D aggregate positions relative to each 2D cell type. **c**, PCA representing each cluster in **b**, designated by WT or KI UDET source.

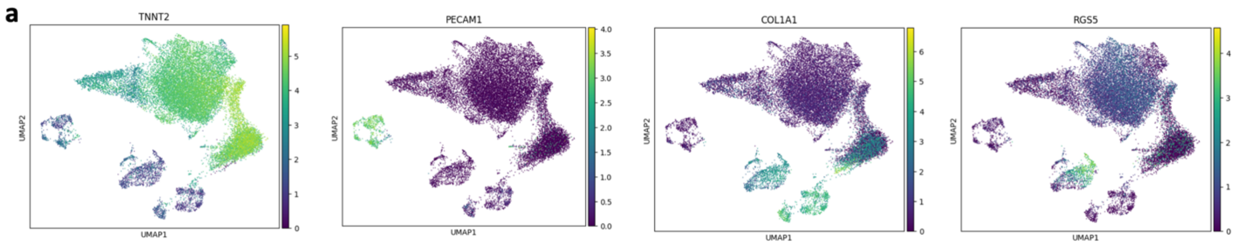


**Supplementary Fig. 13: Generation of annotated UMAPs from cardiac organoid single-cell RNA sequencing data. a**, DimPlots highlighting localization of key annotation markers from cardiac organoids data.

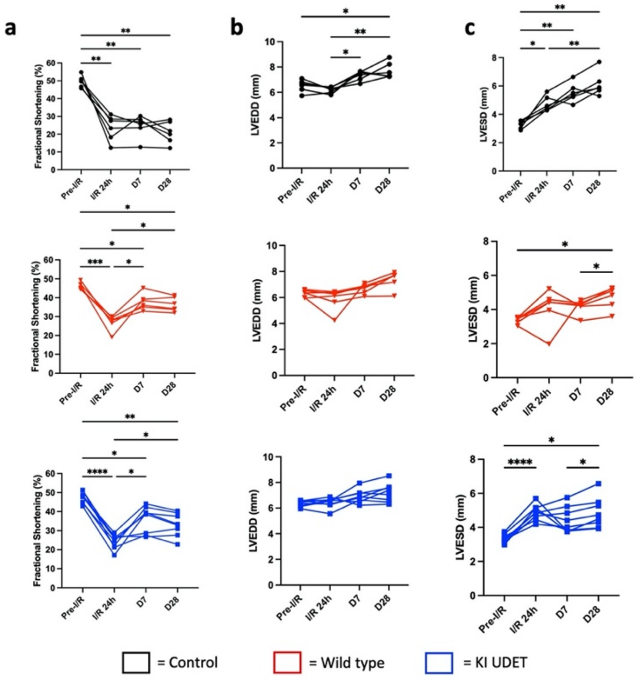


**Supplementary Fig. 14: Echocardiographic values corresponding to each I/R injured rat. a,** FS of each rat at pre-I/R, post-I/R, and 7 and 28 days after transplantation. (n = 6, 6, and 8 rats for control, WT, and KI UDET organoid treatment groups, respectively). **b**, LVEDD of each rat at pre-I/R, post-I/R, and 7 and 28 days after transplantation. **c**, LVESD of each rat at pre-I/R, post-I/R, and 7 and 28 days after transplantation.

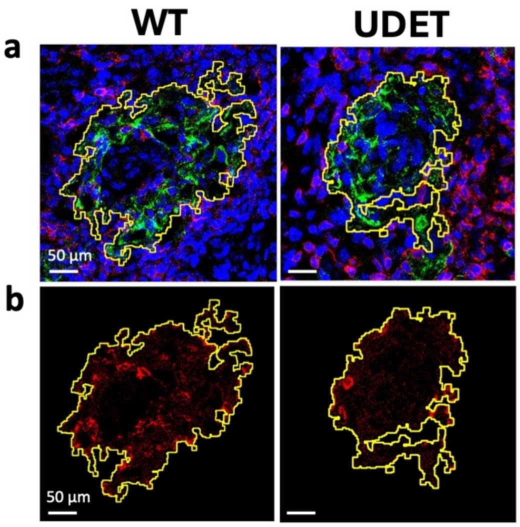


**Supplementary Fig.15: Graft area assessment in cardiac organoid kidney capsule transplantations. a**, Graft area as defined by cTnT+ signal. **b**, CD3+ signal within graft area defined in **a**.

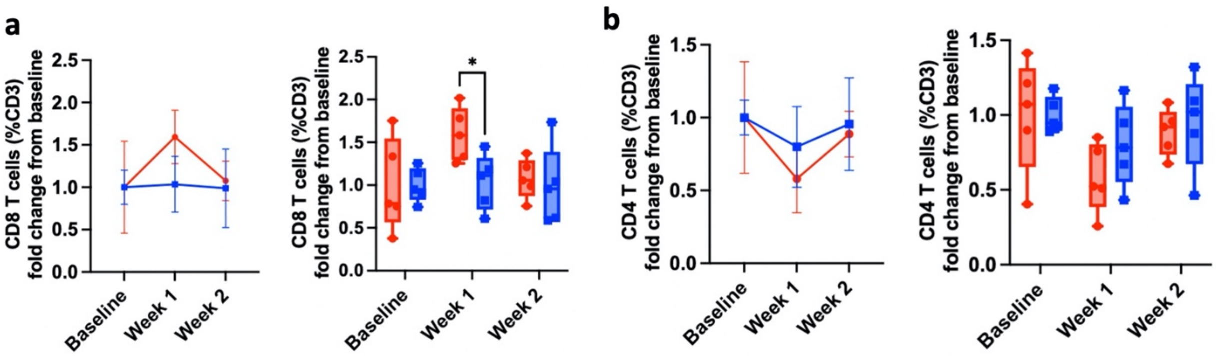


**Supplementary Fig. 16: Circulating CD4+ and CD8+ T cell populations in mice with cardiac organoid kidney capsule transplants. a**, Quantification of flow cytometry data evaluating CD8+ cells in circulation as percent of CD3+ cells in mice receiving cardiac organoid kidney capsule transplantations, displayed as fold change from baseline. **b,** Quantification of flow cytometry data evaluating CD4+ cells in circulation as percent of CD3+ cells in mice receiving cardiac organoid kidney capsule transplantations, displayed as fold change from baseline*=p<0.05.
